# ER-to-lysosome Ca^2+^ refilling followed by K^+^ efflux-coupled store-operated Ca^2+^ entry in inflammasome activation and metabolic inflammation

**DOI:** 10.1101/2023.04.19.537448

**Authors:** Hyereen Kang, Seong Woo Choi, Joo Young Kim, Soo-Jin Oh, Sung Joon Kim, Myung-Shik Lee

**Affiliations:** Severance Biomedical Science Institute, Yonsei University College of Medicine, Seoul 03722, KOREA; Department of Physiology and Ion Channel Disease Research Center, Dongguk University College of Medicine, Gyeongju, Gyeongsangbuk-do 38066, KOREA; Department of Pharmacology and Brain Korea 21 Project for Medical Sciences, Yonsei University College of Medicine, Seoul 03722, KOREA; Soonchunhyang Institute of Medi-bio Science and Division of Endocrinology, Department of Internal Medicine, Soonchunhyang University College of Medicine, Cheonan 31151, KOREA; Department of Physiology, Ischemic/Hypoxic Disease Institute, Seoul National University College of Medicine, Seoul 03080, KOREA

## Abstract

We studied lysosomal Ca^2+^ in inflammasome. LPS+palmitic acid (PA) decreased lysosomal Ca^2+^ ([Ca^2+^]_Lys_) and increased [Ca^2+^]_i_ through mitochondrial ROS, which was suppressed in *Trpm2*-KO macrophages. Inflammasome activation and metabolic inflammation in adipose tissue of high-fat diet (HFD)-fed mice were ameliorated by *Trpm2* KO. ER→lysosome Ca^2+^ refilling occurred after lysosomal Ca^2+^ release whose blockade attenuated LPS+PA-induced inflammasome. Subsequently, store-operated Ca^2+^entry (SOCE) was activated whose inhibition suppressed inflammasome. SOCE was coupled with K^+^ efflux whose inhibition reduced ER Ca^2+^ content ([Ca^2+^]_ER_) and impaired [Ca^2+^]_Lys_ recovery. LPS+PA activated KCa3.1 channel, a Ca^2+^-activated K^+^ channel. Inhibitors of KCa3.1 channel or *Kcnn4* KO reduced [Ca^2+^]_ER_, attenuated increase of [Ca^2+^]_i_ or inflammasome activation by LPS+PA, and ameliorated HFD-induced inflammasome or metabolic inflammation. Lysosomal Ca^2+^ release induced delayed JNK and ASC phosphorylation through CAMKII-ASK1. These results suggest a novel role of lysosomal Ca^2+^ release sustained by ER→lysosome Ca^2+^ refilling and K^+^ efflux through KCa3.1 channel in inflammasome activation and metabolic inflammation.

## Introduction

Lysosomotropic agents are classical inflammasome activators (Hornung et al., 2008; Misawa et al., 2013; Xian et al., 2022). Detailed mechanism of inflammasome activation by lysosomal stress has been unclear, while roles of lysosomal Ca^2+^ release was suggested (Okada et al., 2014). K^+^ efflux is crucial in most inflammasome activations, facilitating NLRP3-NEK7 oligomerization (He et al., 2016; Sharif et al., 2019). . Relationship between of K^+^ efflux and Ca^2+^ flux in inflammasome has hardly been studied, despite their potential link (Yaron et al., 2015).

We studied lysosomal events in inflammasome focusing on roles of lysosomal Ca^2+^ release by lipopolysaccharide (LPS)+palmitic acid (PA), an effector of metabolic stress (Nakamura et al., 2009) (LP). We found that LP induces mitochondrial reactive oxygen species (ROS) that activates a lysosomal Ca^2+^ efflux channel (TRPM2), leading to lysosomal Ca^2+^ release, delayed JNK activation, ASC phosphorylation and inflammasome activation. We also found occurrence of ER→lysosome Ca^2+^ refilling sustaining lysosomal Ca^2+^ efflux and subsequent store-operated Ca^2+^ entry (SOCE) in inflammasome. Finally, we elucidated roles of K^+^ efflux facilitating SOCE through hyperpolarization-accelerated extracellular Ca^2+^ influx (Guéguinou et al., 2014), and identified KCa3.1 Ca^2+^-activated K^+^ channel as the K^+^ efflux channel in LP-induced inflammasome and metabolic inflammation.

## Results

### Lysosomal Ca^2+^ release by mitochondrial ROS in inflammasome

We investigated whether lysosomal Ca^2+^ release occurs in inflammasome activation by LP, a combination activating inflammasome related to metabolic inflammation (Wen et al., 2011).

When we studied perilysosomal Ca^2**+**^ release in bone marrow-derived macrophages (Mφs) (BMDMs) transfected with *GCaMP3-ML1* (Shen et al., 2012), lysosomal Ca^2+^ release was not directly visualized by LP; however, perilysosomal Ca^2+^ release by Gly-Phe β- naphthylamide, (GPN), a lysosomotropic agent (Shen et al., 2012) was significantly reduced (***Figure 1A***), suggesting preemptying or release of lysosomal Ca^2+^ by LP, similar to the results using other inducers of lysosomal Ca^2+^ release (Park et al., 2022; Zhang et al., 2016). We next measured lysosomal Ca^2+^ content ([Ca^2+^]_Lys_) that can be affected by lysosomal Ca^2+^ release. [Ca^2+^]_Lys_ determined using Oregon Green BAPTA-1 Dextran (OGBD) was significantly lowered by LP (***Figure 1B***), consistent with lysosomal Ca^2+^ release. Likely due to lysosomal Ca^2+^ release, [Ca^2+^]_i_ measured by Fluo-3-AM staining was significantly increased by LP (***Figure 1C***). Ratiometric [Ca^2+^]_i_ measurement using Fura-2 to avoid uneven loading or quenching, validated [Ca^2+^]_i_ increase by LP (***Figure 1C***). Functional roles of increased [Ca^2+^]_i_ in inflammasome by LP was revealed by abrogation of IL-1β release or IL- 1β maturation by BAPTA-AM, a cell-permeable Ca^2+^ chelator (***Figure 1D***). Immunoblotting (IB) demonstrated that pro-IL-1β level was not notably affected by BAPTA-AM, suggesting that effect of BAPTA-AM was unrelated to the potential inhibition of pro-IL-1β transcription or translation (***Figure 1D***).

**Figure 1.**
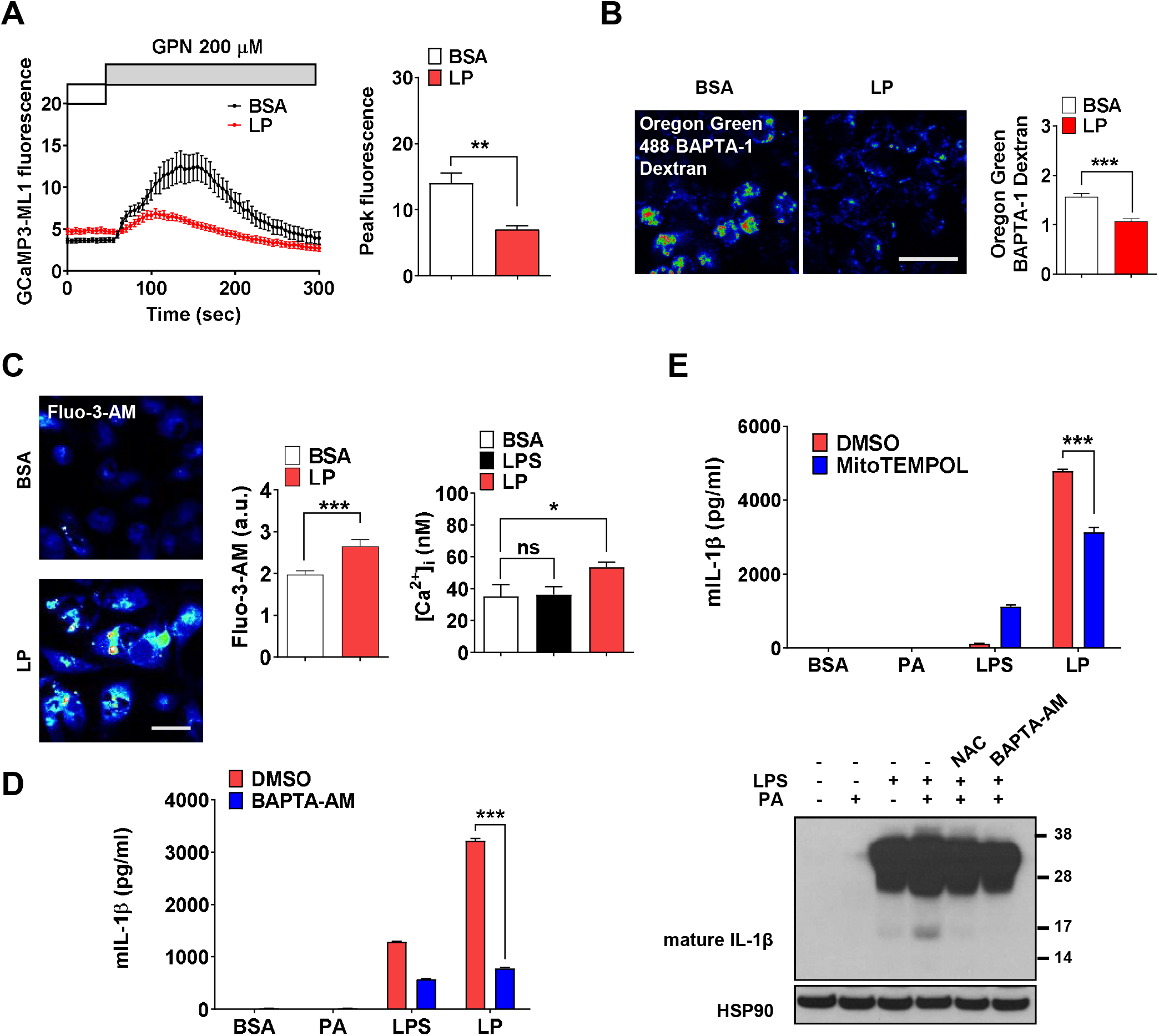
Lysosomal Ca^2+^ and mitochondrial ROS in inflammasome. (**A**) Perilysosomal fluorescence after applying GPN to *GCaMP3-ML1*-transfected BMDMs treated with LP for a total of 4 h including LPS pretreatment for 3 h (left) (actual LP treatment time is 1 h). Peak fluorescence (right) (*n*=6). (**B**) [Ca^2+^]_Lys_ in OGBD-loaded Mφs treated with LP for a total of 4 h including LPS pretreatment for 3 h (right). Representative fluorescence images (left) (*n*=8). (**C**) [Ca^2+^]_i_ in Mφs treated with LP for a total of 4 h including LPS pretreatment for 3 h, determined using Fluo-3-AM staining (middle) or Fura-2 (right). Representative Fluo-3 images (left). (*n*=7 for BSA; *n*=6 for LPS; *n*=13 for LP). (**D**) IL-1β ELISA of culture supernatant after treating Mφs with LPS alone or PA alone for 21 h, or with LP for a total of 21 h including LPS pretreatment for 3 h in the presence or absence of BAPTA-AM (*n*=3) (left). Immunoblotting (IB) of lysate of Mφs treated with LPS alone or PA alone for 21 h, or with LP for a total of 21 h including LPS pretreatment for 3 h in the presence or absence of BAPTA-AM or NAC, using indicated Abs (right). (**E**) IL-1β ELISA of culture supernatant after treating Mφs with LPS alone or PA alone for 21 h, or with LP for a total of 21 h including LPS pretreatment for 3 h in the presence or absence of MitoTEMPOL (*n*=3). Data shown as means ± SEM from more than 3 independent experiments. *p < 0.05, ** p < 0.01 and *** p < 0.001 by two-tailed Student’s *t*-test (A, B), one-way ANOVA with Tukey’s test (C), or two-way ANOVA with Sidak test (D, E) (ns, not significant). Scale bars, 20 μm. The online version of this article includes the following figure supplement for figure 1: **Figure supplement 1.** Mitochondrial ROS in inflammasome activation by LPS+PA (LP).

We next studied mechanism of lysosomal Ca^2+^ release by LP. Since several (lysosomal) ion channels can be activated by ROS (Zhang et al., 2016) and ROS can be produced by PA due to mitochondrial complex inhibition (Nakamura et al., 2009), we studied ROS accumulation. LP effectively induced CM-H2DCFDA fluorescence indicating ROS accumulation, while PA alone or LPS alone induced only a little ROS accumulation (***Figure 1-figure supplement 1A***), suggesting synergistic effect of LPS and PA. When we studied roles of ROS in [Ca^2+^]_i_ increase using *N*-acetyl cysteine (NAC), an antioxidant (Tardiolo et al., 2018), release and maturation of IL-1β by LP were significantly reduced (***Figure 1-figure supplement 1B and Figure 1D***). NAC abrogated LP-induced increase of [Ca^2+^]_i_ as well (***Figure 1-figure supplement 1C***), substantiating functional roles of ROS in [Ca^2+^]_i_ increase and inflammasome activation. We also studied mitochondrial ROS because mitochondria is a well-known target of PA, an effector of metabolic stress (Nakamura et al., 2009) and critical in inflammasome (Xian et al., 2022; Zhou et al., 2011). Using MitoSOX, we observed significant mitochondrial ROS accumulation by LP. When we quenched mitochondrial ROS using MitoTEMPOL (***Figure 1-figure supplement 1D***), IL-1β release by LP was significantly reduced (***Figure 1E***), suggesting crucial roles of mitochondrial ROS in LP- induced inflammasome.

### Ca^2+^ release through lysosomal TRPM2 channel in inflammasome

We next studied which lysosomal Ca^2+^ exit channel is involved in lysosomal Ca^2+^ release by LP. Previous papers suggested roles of transient receptor potential melastatin 2 (TRPM2) channel on the plasma membrane, in inflammasome by other stimulators (Tseng et al., 2017; Wang et al., 2020). We hypothesized that TRPM2 on lysosome could be involved in LP- induced inflammasome since TRPM2 has been reported to be expressed on lysosome as well (Sumoza-Toledo and Penner, 2010) and *Trpm2*-KO mice are resistant to diet-induced glucose intolerance (Zhang et al., 2012). We verified expression of TRPM2 on lysosome of BMDMs by colocalization of TRPM2 and LAMP2 (***Figure 2-figure supplement 1A***). While physiological ligand of TRPM2 is ADP-ribose (ADPR), ROS can activate TRPM2 by increasing ADPR production from poly-ADPR (Wang et al., 2020). We thus studied effects of apigenin or quercetin inhibiting ADPR generation through CD38 inhibition (Bock, 2020; Nam et al., 2020). Inflammasome by LP was significantly inhibited by apigenin or quercetin (***Figure 2-figure supplement 1B***), suggesting that TRPM2 participates in inflammasome by LP. [Ca^2+^]_i_ increase by LP was also reduced by apigenin or quercetin (***Figure 2-figure supplement 1C***), supporting that apigenin or quercetin suppresses inflammasome through inhibition of lysosomal Ca^2+^ channels activated by ADPR such as TRPM2. Since apigenin or quercetin has effects other than CD38 inhibition (Li et al., 2016; Zhang et al., 2020), we employed *Trpm2*-KO mice. [Ca^2+^]_i_ increase and [Ca^2+^]_Lys_ decrease by LP were abrogated in *Trpm2*-KO Mφs (***Figures 2A and B***), suggesting roles of TRPM2 in lysosomal Ca^2+^ release by LP. However, ROS production by LP was not changed by *Trpm2* KO (***Figure 2-figure supplement 1D***), suggesting that TRPM2 is downstream of ROS. Importantly, inflammasome activation assessed by IL-1β ELISA of culture supernatant or IB of Mφ lysate after LP treatment was significantly reduced by *Trpm2* KO (***Figure 2C***), indicating roles of TRPM2, likely lysosomal TRPM2, in inflammasome by LP. ASC speck formation, a marker of inflammasome, by LP was also markedly reduced by *Trpm2* KO (***Figure 2D***). When we employed other activators, inflammasome activation by L-Leucyl-L-Leucine methyl ester (LLOMe) or monosodium urate (MSU), lysosomotropic agents, was significantly inhibited by *Trpm2* KO. However, inflammasome by nigericin, an ionophore exchanging K^+^ for H^+^ or ATP acting on P2X7 ion channel on the plasma membrane (Campden and Zhang, 2019), was not significantly affected (***Figure 2-figure supplement 1E***), suggesting that TRPM2 channel is crucial in inflammasome involving lysosomal Ca^2+^.

**Figure 2.**
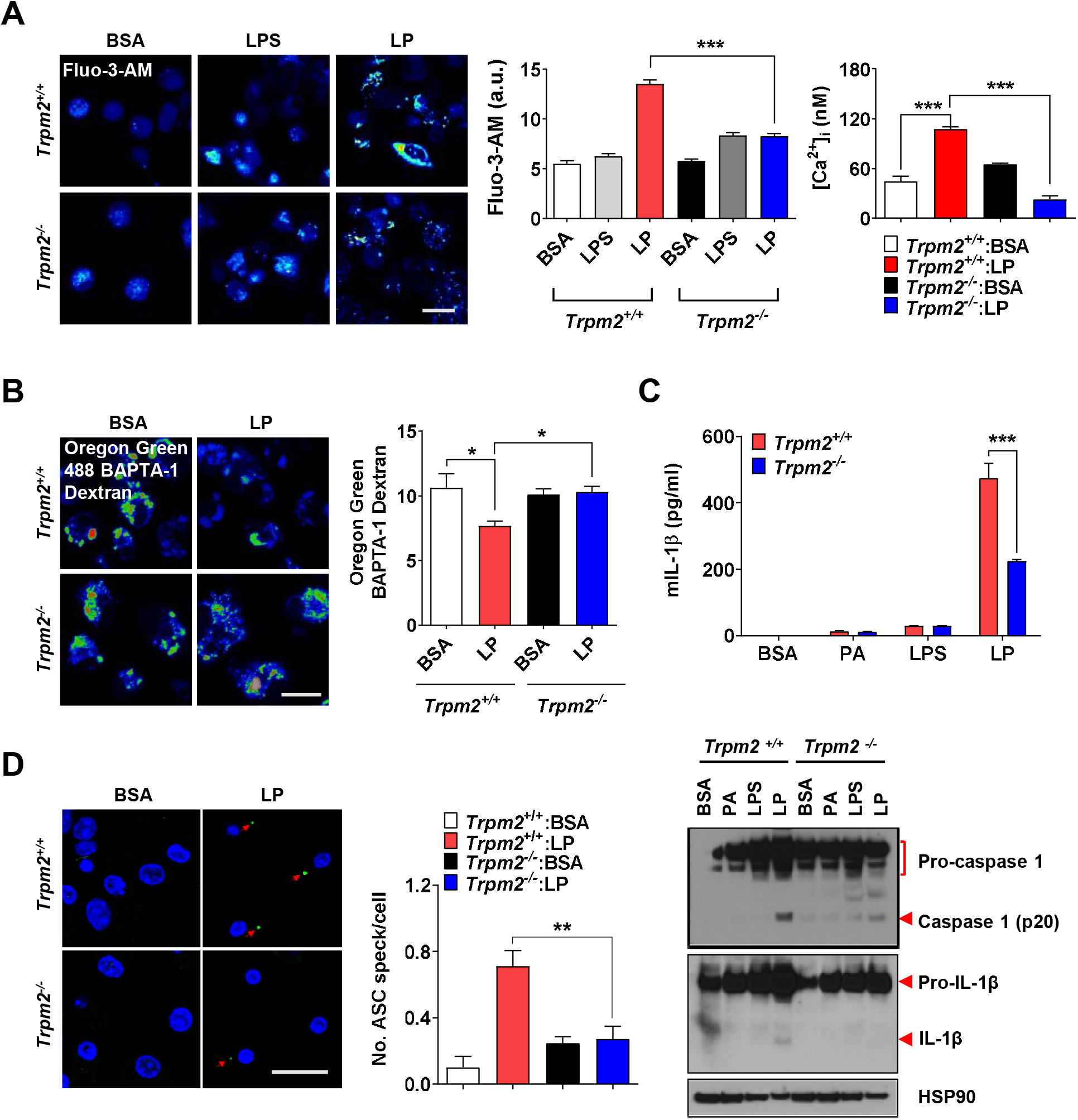
Lysosomal Ca^2+^ efflux through TRPM2 in inflammasome. (**A**) [Ca^2+^]_i_ in Mφs treated with LPS alone for 4 h or with LP for a total of 4 h including LPS pretreatment for 3 h, determined using Fluo-3-AM (middle) or Fura-2 (right). Representative Fluo-3 images (left). (n=8 for Fluo-3-AM; n=9 for Fura-2). (**B**) [Ca^2+^]_Lys_ in Mφs treated with LP for a total of 4 h including LPS pretreatment for 3 h, determined by OGBD loading (right). Representative fluorescence images (left). (n=5) (**C**) IL-1β ELISA of culture supernatant (upper) and IB of cell lysate using indicated Abs after treatment of Mφs with LPS alone for 21 h or LP for a total of 21 h including LPS pretreatment for 3 (lower). (n=4) (**D**) The number of ASC specks in Mφs treated with LP for a total of 21 h including LPS pretreatment for 3 h, determined by immunofluorescence using anti-ASC Ab (right). Representative confocal images (left). (*n*=4 for *Trpm2*^+/+^:BSA; *n*=5 for *Trpm2*^+/+^: LP; *n*=5 for *Trpm2^-^*^/-^:BSA; *n*=7 for *Trpm2^-^*^/-^:LP) Data shown as means ± SEM from more than 3 independent experiments. *p < 0.05, **p < 0.01 and ***p < 0.001 by one-way ANOVA with Tukey’s test (A, B, D), or two-way ANOVA with Sidak test or Bonferroni test (C). Scale bars, 20 μm. The online version of this article includes the following figure supplement for figure 2: **Figure supplement 1.** Effect of CD38 inhibitors and *Trpm2* KO on inflammasome.

Despite its crucial role in inflammasome by LP, TRPM2 exists on both plasma membrane and lysosome. We thus studied whether plasma membrane TRPM2 current can be activated by LP employing *N*-(*p*-amylcinnamoyl)anthranilic acid (ACA), an inhibitor of TRPM2 (Kraft et al., 2006). ACA could inhibit IL-1β release by LP as expected (***Figure 3-figure supplement 1A***). Intracellular dialysis using cyclic ADPR induced slow inward current on the plasma membrane, which was inhibited by ACA (***Figure 3-figure supplement 1B***). Amplitude of cyclic ADPR-induced inward current was not affected by PA and/or LPS (b-c in ***Figure 3-figure supplement 1B and C***). Basal inward current inhibited by ACA, i.e., unstimulated TRPM2 activity, was also not changed by PA and/or LPS (a-c in ***Figure 3*- *figure supplement 1B and C***), suggesting that LP neither affects nor induces plasma membrane TRPM2 current and that lysosomal TRPM2 is likely important in LP-induced inflammasome. To further study roles of lysosomal TRPM2 in inflammasome activation by LP, we employed bafilomycin A1 emptying lysosomal Ca^2+^ reservoir through lysosomal v-ATPase inhibition (Kinnear et al., 2004; Lange et al., 2009). TRPM2-dependent [Ca^2+^]_i_ increase by LP was abrogated by bafilomycin A1 (***Figure 3-figure supplement 1D***), strongly supporting Ca^2+^ release through lysosomal TRPM2 by LP.

**Figure 3.**
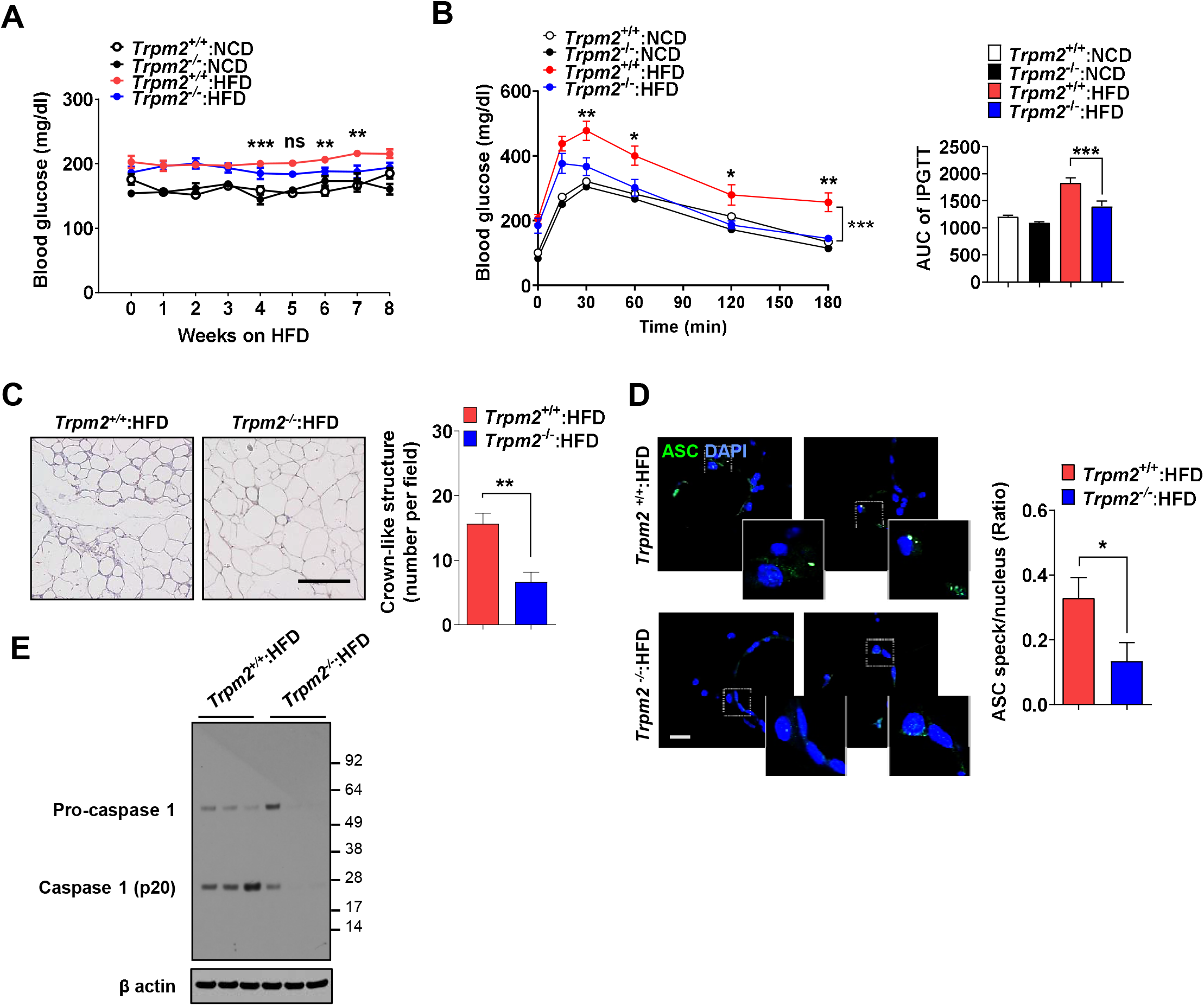
Ameliorated metabolic inflammation by *Trpm2* KO. (**A**) Nonfasting blood glucose of mice on normal chow diet (NCD) (*n*=5 each) or HFD (*n*=8 each). (*: comparison between *Trpm2*^+/+^ and *Trpm2*^-/-^ mice on HFD). (**B**) IPGTT after NCD (*n*=5 each) or HFD (*n*=8 each) feeding for 8 weeks (left). AUC (right). (**C**) The number of CLS in WAT after HFD feeding for 8 weeks (right). Representative H&E sections (left). (scale bar, 50 μm) (*n*=8 each). (**D**) The number of ASC specks in WAT after HFD feeding for 8 weeks, determined by immunofluorescence using anti-ASC Ab (right). Representative confocal images (left). (scale bar, 20 μm) (insets, magnified) (*n*=7 each). (**E**) IB of SVF of WAT after HFD feeding for 8 weeks using indicated Abs. (*n*=3). Data shown as means ± SEM from more than 3 independent experiments. *p < 0.05, **p < 0.01 and ***p < 0.001 by two-way ANOVA (B) or two-tailed Student’s *t*-test (A, C, D). The online version of this article includes the following figure supplement for figure 3: **Figure supplement 1.** Plasma membrane TRPM2 current in Mφs and metabolic profile of *Trpm2*-KO mice.

### Ameliorated metabolic inflammation by *Trpm2* KO

Since lysosomal TRPM2 is likely involved in inflammasome by LP, we next studied roles of TRPM2 in inflammasome and metabolic inflammation in vivo. Nonfasting blood glucose was significantly lower in *Trpm2*-KO mice on HFD compared to control mice on HFD (***Figure 3A***), while body weight was not different between them (***Figure 3-figure supplement 1E***). Intraperitoneal glucose tolerance test (IPGTT) showed significantly ameliorated glucose intolerance and reduced area under the curve (AUC) in *Trpm2*-KO mice on HFD (***Figure 3B***). HOMA-IR, an index of insulin resistance, was also significantly reduced in *Trpm2*-KO mice

### on HFD (Figure 3-figure supplement 1F)

We studied whether improved metabolic profile by *Trpm2* KO is due to reduced inflammasome. The number of crown-like structures (CLS) representing metabolic inflammation (Weisberg et al., 2003) was significantly reduced in while adipose tissue (WAT) of *Trpm2*-KO mice on HFD (***Figure 3C***), suggesting reduced metabolic inflammation by *Trpm2* KO. Inflammasome activation was also significantly reduced in WAT of *Trpm2*-KO mice on HFD as evidenced by significantly reduced numbers of ASC specks and capase-1 cleavage (***Figures 3D and E***), indicating that TRPM2 is important in inflammasome related to metabolic syndrome.

### ER→lysosome Ca^2+^ refilling in inflammasome

After confirming roles of lysosomal TRPM2 in [Ca^2+^]_i_ increase by LP, we studied changes of ER Ca^2+^ content ([Ca^2+^]_ER_) as ER, the largest intracellular Ca^2+^ reservoir, interacts with other organelles in cellular processes requiring intracellular Ca^2+^ flux. While [Ca^2+^]_Lys_ is comparable to [Ca^2+^]_ER_ (Raffaello et al., 2016), lysosome alone might not be a major Ca^2+^ source because of small volume (Penny et al., 2014). Thus, we studied whether ER to lysosomal Ca^2+^ flux which has been observed after lysosomal Ca^2+^ emptying (Garrity et al., 2016; Park et al., 2022), occurs during inflammasome activation to sustain lysosomal Ca^2+^ release. When we measured [Ca^2+^]_ER_ in *GEM-CEPIA1er* (Suzuki et al., 2014)-transfected BMDMs treated with LP without extracellular Ca^2+^ to abolish SOCE (Derler et al., 2016), [Ca^2+^]_ER_ became significantly lower (***Figure 4A***), suggesting Ca^2+^ flux from ER to lysosome likely to replenish reduced [Ca^2+^]_Lys_ during inflammasome. [Ca^2+^]_ER_ determined using a FRET-based *D1ER* (Park et al., 2009) also demonstrated a significantly reduced [Ca^2+^]_ER_ by LP in a Ca^2+^-free KRB buffer (***Figure 4A***). To study dynamic changes of [Ca^2+^]_ER_ and its temporal relationship with [Ca^2+^]_Lys_, we simultaneously traced [Ca^2+^]_ER_ and [Ca^2+^]_Lys_ in *GEM- CEPIA1er*-transfected cells loaded with OGBD. When [Ca^2+^] was monitored in cells that have reduced [Ca^2+^]_Lys_ after LP treatment and then were incubated without LP in a Ca^2+^-free medium, recovery of decreased [Ca^2+^]_Lys_ was noted (***Figure 4B***). In this condition, [Ca^2+^]_ER_ decrease occurred in parallel with [Ca^2+^]_Lys_ recovery (***Figure 4B***), strongly indicating ER→lysosome Ca^2+^ refilling.

**Figure 4.**
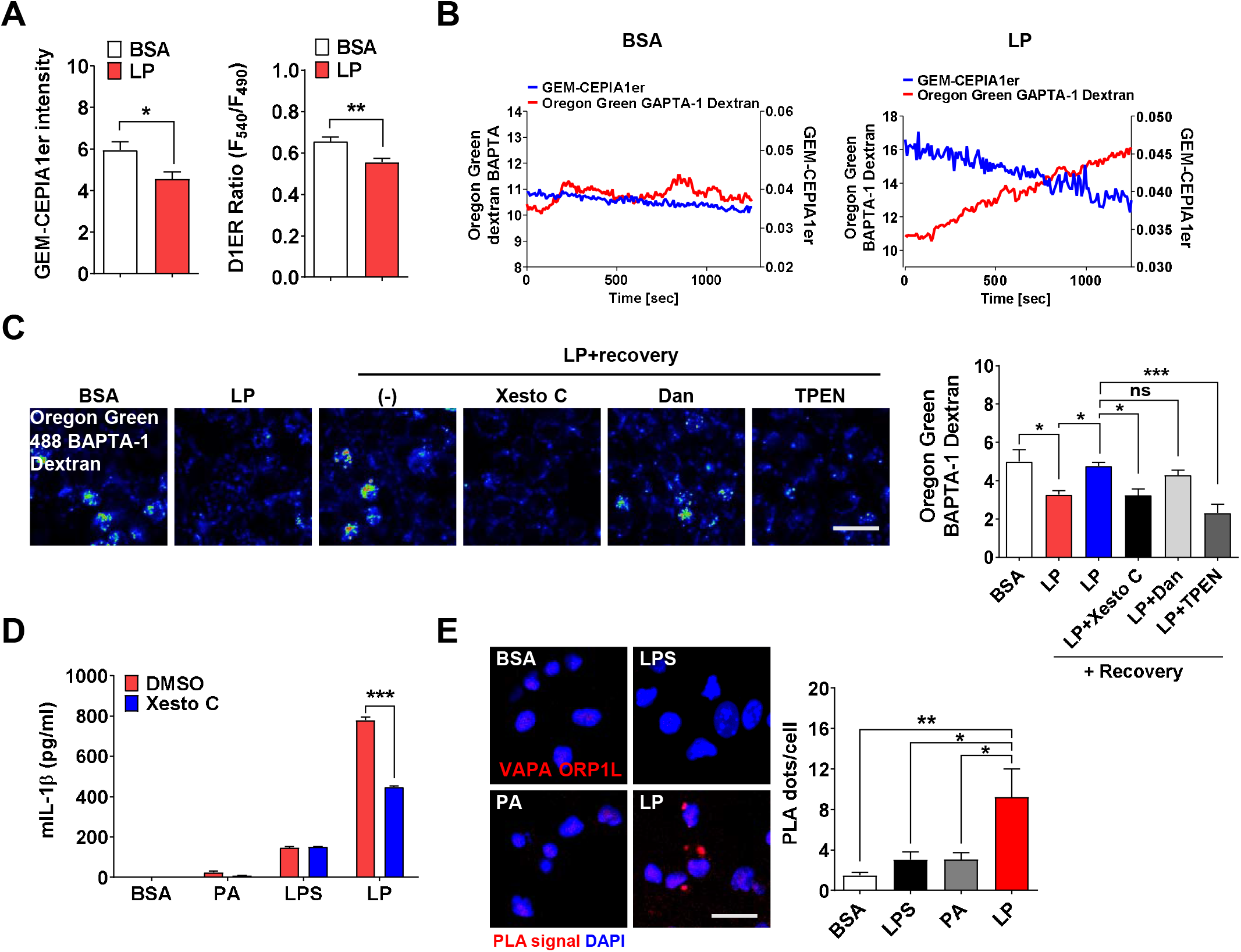
ER→lysosome Ca^2+^ refilling in inflammasome. (**A**) [Ca^2+^]_ER_ in *GEM-CEPIA1e*r- (left) or *D1ER-*transfected BMDMs (right) treated with LP for 1 h without extracellular Ca ^2+^ after LPS pretreatment for 3 h. (*n*=26 for BSA; *n*=25 for LP). (**B**) BMDMs transfected with *GEM-CEPIA1er* and loaded with OGBD were treated with LP for 1 h after LPS pretreatment for 3 h (right) or BSA alone for 4 h (left). Tracing of [Ca^2+^]_Lys_ and [Ca^2+^]_ER_ after change to a fresh medium without extracellular Ca^2+^. (*n*=4 for BSA; *n*=4 for LP). (**C**) OGBD-loaded Mφs were treated with LP for 1 h after LPS pretreatment for 3 h. Recovery of [Ca^2+^]_Lys_ after change to a fresh medium with or without Xestospongin C (Xesto C), dantrolene (Dan) or TPEN (right). Representative confocal images (left) (*n*=9 for BSA; *n*=8 for LP; *n*=9 for LP+Recovery; *n*=6 for LP+Recovery+Xesto C; *n*=5 for LP+Recovery+Dan; *n*=8 for LP+Recovery+TPEN). (**D**) IL-1β ELISA of culture supernatant after treating Mφs with LPS alone or PA alone for 21 h, or with LP for a total of 21 h including LPS pretreatment for 3 h, in the presence or absence of Xesto C (*n*=4). (**E**) PLA in Mφs treated with LPS alone or PA alone for 21 h, or LP for a total of 21 h including LPS pretreatment for 3 h, using Abs to VAPA and ORP1L (right). Representative fluorescence images (left) (n=4). Data shown as means ± SEM from more than 3 independent experiments. *p < 0.05, **p < 0.01 and ***p < 0.001 by two-tailed Student’s t-test (A), one-way ANOVA with Tukey’s test (C, E) or two- way ANOVA with Sidak test (D). Scale bar, 20 μm. The online version of this article includes the following figure supplement for figure 4: **Figure supplement 1.** Store-operated Ca^2+^ entry (SOCE) in inflammasome by LP.

We next studied which ER Ca^2+^ exit channels are involved in ER→lysosome Ca^2+^ refilling. When LP was removed after treatment, [Ca^2+^]_Lys_ recovery was observed in a Ca^2+^- replete medium (***Figure 4C***). Here, Xestospongin C, an IP_3_ receptor (IP_3_R) channel antagonist (Garrity et al., 2016), inhibited recovery of [Ca^2+^]_Lys_ after LP removal (***Figure 4C***), suggesting ER→lysosome Ca^2+^ refilling through IP_3_R channel. Dantrolene, an antagonist of ryanodine receptor (RyR) channel, another ER Ca^2+^ exit channel (Garrity et al., 2016), did not significantly affect recovery of [Ca^2+^]_Lys_ (***Figure 4C***). When we chelated ER Ca^2+^ with a membrane-permeant metal chelator *N,N,N’,N’*-tetrakis (2-pyridylmethyl)ethylene diamine (TPEN) that has a low Ca^2+^ affinity and can chelate ER Ca^2+^ but not cytosolic Ca^2+^ (Hofer et al., 1998), recovery of [Ca^2+^]_Lys_ after LP removal was markedly inhibited (***Figure 4C***), again supporting ER→lysosome Ca^2+^ refilling during lysosomal Ca^2^ recovery. When functional impact of ER→lysosome Ca^2+^ refilling was studied, Xestospongin C significantly suppressed IL-1β release by LP (***Figure 4D***), suggesting that ER→lysosome Ca^2+^ refilling contributes to inflammasome by LP. Since ER→lysosome Ca^2+^ refilling could be facilitated by membrane contact between organelles (Yang et al., 2019), we studied apposition of ER and lysosome proteins. Proximity ligation assay (PLA) demonstrated multiple contact between VAPA on ER and ORP1L on lysosome by LP (***Figure 4E***), suggesting facilitation of ER→lysosome Ca^2+^ refilling by organelle contact.

As extracellular Ca^2+^ is likely to enter cells through SOCE channel after ER Ca^2+^ depletion (Derler et al., 2016), we next studied extracellular Ca^2+^. When extracellular Ca^2+^ was chelated by 3 mM EGTA reducing [Ca^2+^]_i_ in RPMI medium to 99 nM (Schoenmakers et al., 1992) below [Ca^2+^]_i_ in Ca^2+^-free medium (Maggi et al., 1989), IL-1β release by LP was significantly reduced (***Figure 4-figure supplement 1A***), demonstrating roles of extracellular Ca^2+^ in inflammasome by LP. 2-APB, a SOCE inhibitor, also significantly reduced IL-1β release by LP (***Figure 4-figure supplement 1B***), supporting roles of SOCE in inflammasome. Since 2-APB can inhibit ER Ca^2+^ channel as well, we next employed another SOCE inhibitor. BTP2, a SOCE inhibitor that does not affect ER Ca^2+^ channel (Zitt et al., 2004), significantly reduced IL-1β release by LP (***Figure 4-figure supplement 1C***), substantiating roles of SOCE in inflammasome. We also studied whether BTP2 can affect [Ca^2+^]_ER_ in LP-induced inflammasome through SOCE inhibition. We determined [Ca^2+^]_ER_ without extracellular Ca^2+^ removal because BTP2 effect on SOCE cannot be seen after extracellular Ca^2+^ removal abrogating SOCE. In the presence of extracellular Ca^2+^, [Ca^2+^]_ER_ was not decreased by LP, likely due to SOCE (***Figure 4-figure supplement 1D***). Here, BTP2 significantly reduced [Ca^2+^]_ER_ of LP-treated BMDMs (***Figure 4-figure supplement 1D***) likely due to SOCE inhibition, suggesting that SOCE is activated in inflammasome by LP to replenish reduced ER Ca^2+^ store. We also studied aggregation of STIM1, a Ca^2+^ sensor in ER, which can be observed in SOCE activation (Derler et al., 2016). Indeed, STIM1 aggregation was clearly observed after LP treatment (***Figure 4-figure supplement 1E***), which was colocalized with ORAI1, an SOCE channel on the plasma membrane (Vaca, 2010), strongly indicating SOCE through ORAI1 channel in LP-induced inflammasome.

### Coupling of K^+^ efflux and Ca^2+^ influx in inflammasome

K^+^ efflux is one of the most common and critical events in inflammasome (Muñoz-Planillo et al., 2013), although a couple of inflammasomes without K^+^ efflux have been reported (Groß et al., 2016). In excitable cells, K^+^ efflux leads to hyperpolarization, which negatively modulates Ca^2+^ influx through voltage-gated Ca^2+^ channel activated by depolarization. In nonexcitable cells, contrarily, Ca^2+^ influx though voltage-independent Ca^2+^ channel such as SOCE channel can be positively modulated by K^+^ efflux, due to increased electrical Ca^2+^ driving force (Guéguinou et al., 2014). Since Ca^2+^ influx from extracellular space into cytosol might be positively modulated by K^+^ efflux in nonexcitable cells such as Mφs, we studied whether K^+^ efflux is coupled to Ca^2+^ influx. Intracellular K^+^ content ([K^+^]_i_) determined using Potassium Green-2-AM was decreased by LP (***Figure 5-figure supplement 1A***), likely due to K^+^ efflux. High extracellular K^+^ content ([K^+^]_e_) (60 mM) inhibited inflammasome activation by LP likely by inhibiting K^+^ efflux (***Figure 5-figure supplement 1B***). We then studied effects of high [K^+^]_e_ on [Ca^2+^]_ER_ that would be affected by high [K^+^]_e_ if K^+^ efflux and Ca^2+^ influx through SOCE are coupled. We determined [Ca^2+^]_ER_ again without extracellular Ca^2+^ removal since possible effects of high [K^+^]_e_ on SOCE cannot be seen after extracellular Ca^2+^ removal. At [K^+^]_e_ of 5.4 mM ([K^+^]_e_ in RPMI), [Ca^2+^]_ER_ was not decreased by LP. However, at high [K^+^]_e_, LP significantly reduced [Ca^2+^]_ER_ (***Figure 5A***) likely due to SOCE inhibition by high [K^+^]_e_, suggesting that K^+^ efflux is coupled to extracellular Ca^2+^ influx through SOCE. We next studied whether [Ca^2+^]_Lys_ recovery which is seen after LP removal due to ER→lysosome Ca^2+^ refilling is affected by high [K^+^]_e_. [Ca^2+^]_Lys_ recovery after LP removal became significantly lower by high [K^+^]_e_ (***Figure 5B***), suggesting that high [K^+^]_e_ dampens [Ca^2+^]_Lys_ recovery after LP removal likely through SOCE inhibition. High [K^+^]_e_ also suppressed [Ca^2+^]_i_ increase by LP (***Figure 5C***), suggesting contribution of K^+^ efflux in [Ca^2+^]_i_ increase by LP through sustained Ca^2+^ influx via SOCE.

**Figure 5.**
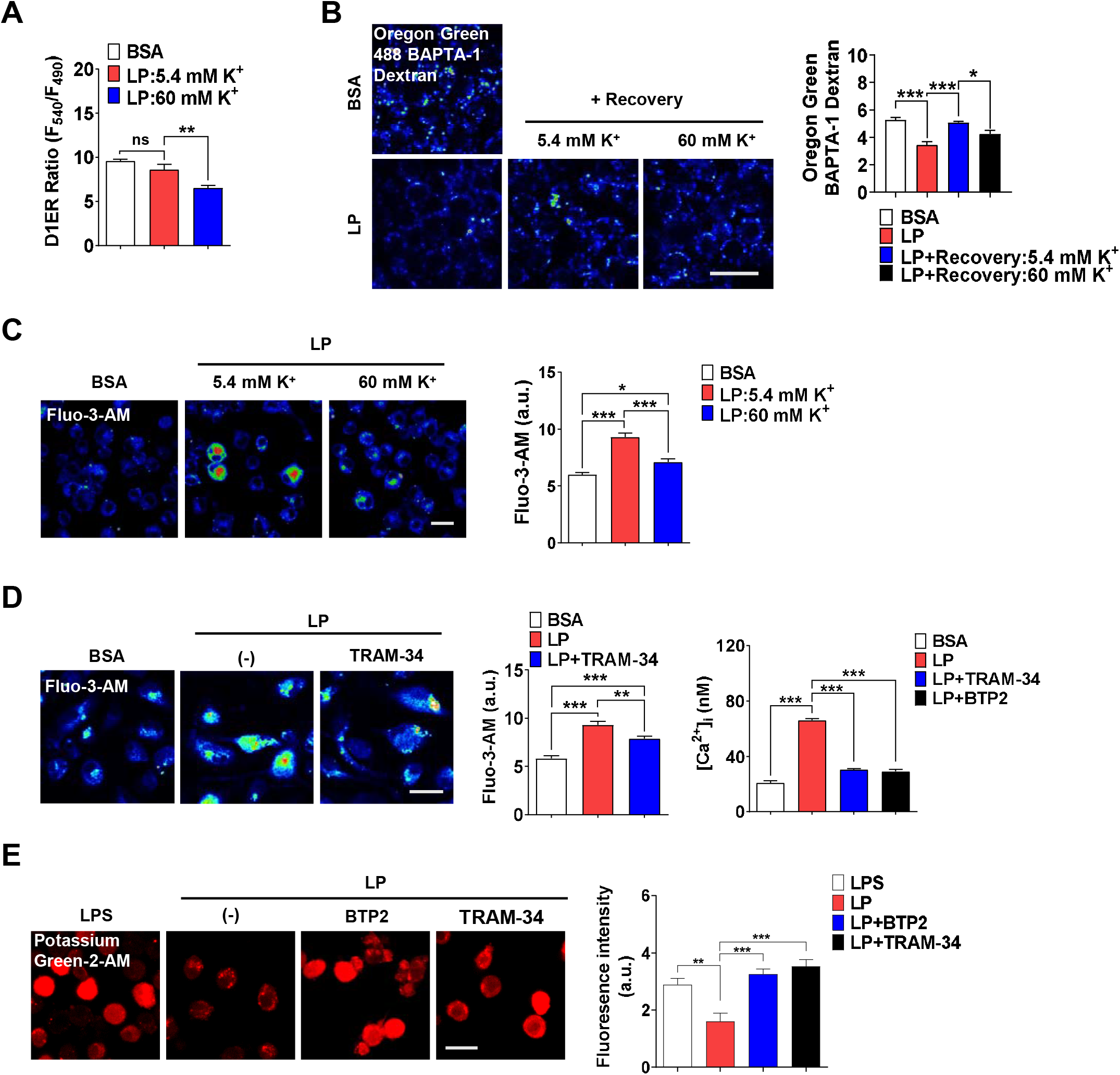
Coupling of K^+^ efflux and Ca^2+^ influx in inflammasome. (**A**) [Ca^2+^]_ER_ in *D1ER-* transfected BMDMs treated with LP for a total of 4 h including LPS pretreatment for 3 h at [K^+^]_e_ of 5.4 or 60 mM (*n*=21 each). (**B**) OGBD-loaded Mφs were treated with LP for a total of 4 h including LPS pretreatment for 3 h. Recovery of [Ca^2+^]_Lys_ in a fresh medium with 5.4 or 60 mM K^+^ (right). Representative fluorescence images (left) (*n*=7 for BSA; *n*=6 for LP; *n*=7 for LP+Recovery:5.4 mM K^+^; *n*=6 for LP+Recovery:60 mM K^+^). (**C**) [Ca^2+^]_i_ in Mφs treated with LP for a total of 4 h including after LPS pretreatment for 3 h in a medium with 5.4 or 60 mM K^+^ (right). Representative Fluo-3 images (left) (*n*=14 for BSA; *n*=8 for LP:5.4 mM K^+^; *n*=6 for LP:60 mM K^+^). (**D**) [Ca^2+^]_i_ in Mφs treated with LP for 1 h in the presence or absence of BTP2 or TRAM-34 after LPS pretreatment for 3 h, determined using Fluo-3-AM (middle) or Fura-2 (right). Representative Fluo-3 images (left) (For Fluo-3-AM, *n*=8) (For Fura-2, *n*=13). (**E**) [K^+^]_i_ after LP treatment for a total of 21 h including LPS pretreatment for 3 h in the presence or absence of BTP2 or TRAM-34 (right). Representative Potassium Green-2 images (left) (*n*=5). Data shown as means ± SEM from more than 3 independent experiments. *p < 0.05, ** p< 0.01 and ***p < 0.001 by one-way ANOVA with Tukey’s test (A-E). Scale bar, 20 μm. The online version of this article includes the following figure supplement for figure 5: **Figure supplement 1.** Effect of high extracellular K^+^ and inhibitors of K^+^ efflux channels on inflammasome.

We next investigated which K^+^ efflux channel is involved in inflammasome, focusing on Ca^2+^-activated K^+^ channels that can positively modulate Ca^2+^ influx after initial Ca^2+^ entry (Guéguinou et al., 2014). When Mφs were incubated with various K^+^ efflux channel inhibitors, charybdotoxin (CTX) inhibiting all 3 types of Ca^2+^-activated K^+^ channels [BK, IKCa1 (KCa3.1) and SK] (Chen and Chung, 2013; González et al., 2012), significantly suppressed IL-1β release by LP (***Figure 5-figure supplement 1C***), supporting roles of Ca^2+^- activated K^+^ channels in inflammasome. In contrast, inhibitors of K^+^ efflux channels unrelated to Ca^2+^-induced activation such as quinine (a 2-pore K^+^ channel inhibitor), barium sulfate (Kir channel inhibitor) or 4-aminopyridine (4-AP, a Kv channel inhibitor) did not significantly inhibit IL-1β release by LP (***Figure 5-figure supplement 1D***). We next studied which channels among Ca^2+^-activated K^+^ channels are involved. Paxillin, a BK channel inhibitor (González et al., 2012) and UCL 1684 or apamin, SK channel inhibitors (Chen et al., 2021; Strøbæk et al., 2000) did not significantly affect IL-1β release by LP (***Figure 5-figure supplement 1D***). In contrast, TRAM-34, a specific IKCa1 (KCa3.1) channel inhibitor (Agarwal et al., 2014; Wulff et al., 2000), significantly decreased IL-1β release by LP (***Figure 5-figure supplement 1C and D***), suggesting involvement of KCa3.1. We next studied effects of TRAM-34 on [Ca^2+^]_ER_. While [Ca^2+^]_ER_ decrease by LP was not seen without extracellular Ca^2+^ removal likely due to SOCE, it was clearly observed when TRAM-34 was added (***Figure 5-figure supplement 1E***), consistent with positive roles of KCa3.1 in Ca^2+^ influx after ER Ca^2+^ emptying. [Ca^2+^]_i_ increase by LP was also inhibited by TRAM-34 (***Figure 5D***), demonstrating contribution of KCa3.1 channel in the increase of [Ca^2+^]_i_. Further, TRAM-34 abrogated decrease of [K^+^]_i_ by LP (***Figure 5E***), indicating roles of KCa3.1 in K^+^ efflux during inflammasome activation. BTP2 also suppressed increase of [Ca^2+^]_i_ and decrease of [K^+^]_i_ by LP (***Figures 5D and E***), likely by inhibiting Ca^2+^ influx that can activate Ca^2+^-activated K^+^ channel such as KCa3.1.

We next studied whether KCa3.1 K^+^ current is activated by LP. If Ca^2+^-activated K^+^ channel coupled to Ca^2+^ influx mediates K^+^ efflux in inflammasome activation, conventional whole-cell patch clamp using Ca^2+^-clamped solution would nullify effects of Ca^2+^ influx, rendering study of increased Ca^2+^ influx impossible. Hence, we employed nystatin-perforated patch clamp technique that leaves elevated [Ca^2+^]_i_ intact, as nystatin pores are permeable to monovalent but not to divalent ions (Akaike and Harata, 1994). When ramp-like voltage clamp was applied to induce brief current-voltage (I/V) curves and TRAM-34-inhibitable current was obtained by digital subtraction, slope of I/V curves sensitive to TRAM-34 was increased by LP but not by LPS alone (***Figures 6A and B***), suggesting increased KCa3.1 activity by LP. Expression of *Kcnn4* was not significantly affected by LPS and/or PA (***Figure 6-figure supplement 1A***), suggesting that increased KCa3.1 activity by LP is not due to *Kcnn4* induction. When we employed *Kcnn4-*KO Mφs that do not express *Kcnn4* (***Figure 6*- *figure supplement 1B***), TRAM-34-inhibitable K^+^ current was not observed before or after LP treatment (***Figures 6C and D***), confirming reliability of results obtained by nystatin-perforated patch clamp technique.

**Figure 6.**
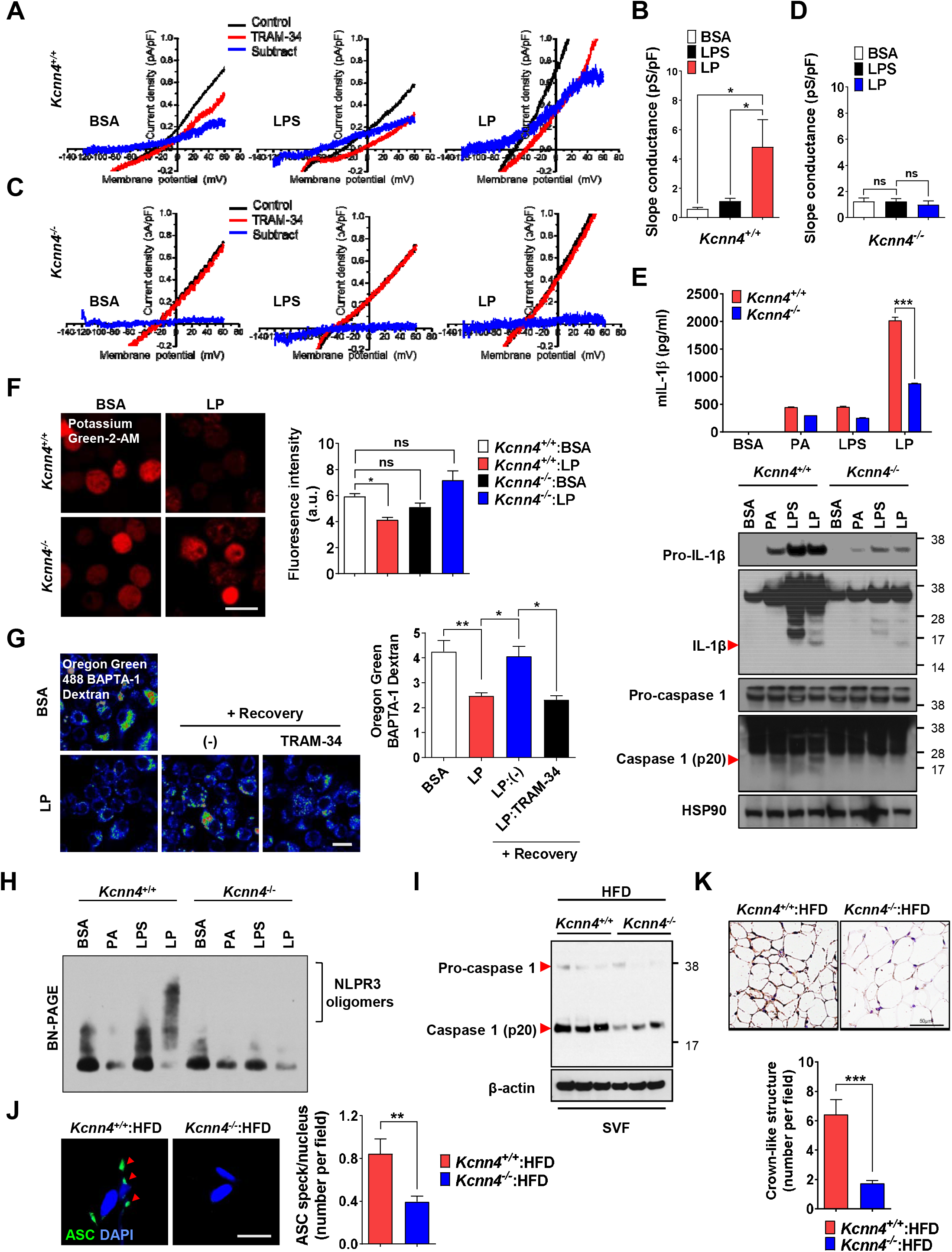
Role of KCa3.1 Ca^2+^-activated K^+^ channel in inflammasome. (**A-D**) Nystatin- perforated patch clamp and slope conductance of TRAM-34-sensitive I/V curve in *Kcnn4*^+/+^ (A, B) or *Kcnn4^-^*^/-^ Mφs (C, D) treated with LPS alone for 4 h or with LP for a total of 4 h including LPS pretreatment for 3 h (B, D). Representative I/V curves (A, C). (*n*=13 for each *Kcnn4*^+/+^ group; *n*=4 for *Kcnn4^-^*^/-^ :BSA; *n*=5 for *Kcnn4^-^*^/-^ :LPS; *n*=4 for *Kcnn4^-^*^/-^ :LP). (**E**) IL- 1β ELISA of culture supernatant (upper) and IB of cell lysate using indicated Abs (lower) after treating *Kcnn4*^+/+^ or *Kcnn4^-^*^/-^ Mφs with PA alone or LPS alone for 21 h, or LP for a total of 21 h including LPS pretreatment for 3 h. (*n*=3). (**F**) [K^+^]_i_ in *Kcnn4*^+/+^ or *Kcnn4^-^*^/-^ Mφs treated with LP for a total of 21 h including LPS pretreatment for 3 h (right). Representative Potassium Green-2 images (left). (*n*=5). (**G**) OGBD-loaded BMDMs were treated with LP for a total of 4 h including LPS pretreatment for 3 h. [Ca^2+^]_Lys_ recovery after LP removal with or without TRAM-34 (right). Representative fluorescence images (left). (*n*=11 for BSA; *n*=8 for LP; *n*=6 for LP:(-); *n*=9 for LP:TRAM-34). (**H**) BN gel electrophoresis and subsequent IB of lysate of Mφs treated with LPS alone for 21 h or with LP for a total of 21 h including LPS pretreatment for 3 h using indicated Ab. (**I**) IB of SVF of WAT from mice fed HFD for 8 weeks using indicated Abs. (*n*=3). (**J**) The number of ASC specks in WAT of mice of (I) identified by ASC immunofluorescence (right). Representative ASC specks (red arrow heads) (left). (scale bar, 20 μm) (*n*=28 for *Kcnn4*^+/+^ :HFD; *n*=22 for *Kcnn4^-^*^/-^ :HFD). (**K**) The number of CLS in WAT of mice of (I) identified by F4/80 immunohistochemistry (lower). Representative F4/80 immunohistochemistry (upper). (scale bar, 50 μm) (*n*=12 for *Kcnn4*^+/+^ :HFD; *n*=11 for *Kcnn4^-^*^/-^ :HFD). Data shown as means ± SEM from more than 3 independent experiments. *p < 0.05, **p < 0.01 and ***p < 0.001 by one-way ANOVA with Tukey’s test (B, D, F, G), or two-way ANOVA with Sidak test (E). Scale bar, 20 μm. The online version of this article includes the following figure supplement for figure 6: **Figure supplement 1.** Effects of *Kcnn4* KO on metabolic profile and inflammasome.

We next studied functional roles of *Kcnn4* in inflammasome. IL-1β release and inflammasome activation by LP were significantly reduced by *Kcnn4* KO (***Figure 6E***), supporting that KCa3.1 channel is important in LP-induced inflammasome. TNF-α or IL-6 release by LPS or LP was not significantly changed by *Kcnn4* KO (***Figure 6-figure supplement 1C***). We also studied effects of *Kcnn4* on the changes of intracellular K^+^ and Ca^2+^ during inflammasome. [K^+^]_i_ decrease by LP was abrogated by *Kcnn4* KO (***Figure 6F***), consistent with roles of KCa3.1 channel in K^+^ efflux. Further, [Ca^2+^]_Lys_ recovery by LP removal, was inhibited by TRAM-34 (***Figure 6G***), indicating that K^+^ efflux contributes to [Ca^2+^]_Lys_ recovery, likely through facilitation of SOCE and subsequent ER→lysosome Ca^2+^ refilling. We also studied physical coupling between Ca^2+^ influx and K^+^ efflux channels, in addition to their functional coupling. PLA demonstrated physical association between KCNN4 and ORAI1 channel by LP (***Figure 6-figure supplement 1D***), which might facilitate functional coupling between K^+^ efflux and SOCE (Ferreira and Schlichter, 2013). Coupling between KCNN4 and ORAI1 was abrogated by BAPTA-AM (***Figure 6-figure supplement 1D***), suggesting roles of increased Ca^2+^ in their directional movement and contact.

We also studied roles of KCa3.1 channel in inflammasome by other stimulators. Inflammasome by LLOMe or MSU, lysosomotropic agents, was significantly inhibited by *Kcnn4* KO, however, that by nigericin directly promoting K^+^ efflux or ATP inducing K^+^ efflux through pannexin-1 channel (Xu et al., 2020; Yang et al., 2015) was not significantly affected (***Figure 6-figure supplement 1E***), suggesting that KCa3.1 channel is crucial in K^+^ efflux associated with lysosomotropic agents or lysosomal Ca^2+^ channels. Since K^+^ efflux is crucial in NLRP3 binding to NEK7 and oligomerization (He et al., 2016), we studied roles of KCa3.1 channel in NLRP3 oligomerization. NLPR3 oligomerization by LP was abrogated by *Kcnn4* KO (***Figure 6H***). NLRP3 binding to NEK7 (He et al., 2016), which was observed in control Mφs treated with LP by immunoprecipitation, was also abrogated by *Kcnn4* KO or TRAM-34 (***Figure 6-figure supplement 1F and G***), suggesting roles of KCa3.1 channel in NLRP3 interaction with NEK7 and oligomerization.

We next studied whether *Kcnn4*-KO mice are resistant to HFD-induced metabolic inflammation. Consistent with roles of KCa3.1 channel in inflammasome by LP, *Kcnn4*-KO mice on HFD showed significantly improved glucose tolerance (***Figure 6-figure supplement 1H***). Inflammasome and metabolic inflammation by HFD were also significantly ameliorated by *Kcnn4* KO as evidenced by attenuated caspase-1 cleavage and significantly reduced number of ASC specks or CLS in WAT (***Figure 6I-K***).

### Mechanism of Ca^2+^-induced inflammasome activation

We next investigated mechanism of inflammasome by increased [Ca^2+^]_i_. We studied whether TAK1-JNK activation observed in inflammasome by lysosomal rupture releasing lysosomal Ca^2+^ (Okada et al., 2014), participates in inflammasome by LP. JNK was activated by LP treatment for 21 h (Figure 7A). Since JNK activation occurs after LPS treatment alone for a short time and then wanes (An et al., 2018), we studied time sequence of JNK activation by LP. JNK activation by LPS alone occurred 30 min after treatment, subsided since 2 h and never occurred again (***Figure 7A***). In contrast, JNK activation occurred again after LP treatment for 21 h (***Figure 7A***), suggesting different mechanism and time scale of JNK activation depending on additional events such as lysosomal ones. While we determined [Ca^2+^]_i_ after LP treatment for a total 4 h including LPS pretreatment for 3 h (actual LP treatment time is 1 h) throughout the study as Ca^2+^ flux is expected to occur since early time, [Ca^2+^]_i_ further increased 21 h after LP treatment (***Figure 7-figure supplement 1A***), which could be high enough to act as signal 2 of inflammasome. [K^+^]_i_ decrease was also observed not 4 but 21 h after LP treatment (***Figure 7-figure supplement 1A***), suggesting that Ca^2+^ influx supported by K^+^ efflux is pronounced at later time. Additionally, [Ca^2+^]_ER_ was reduced not 4 but 21 h after LP treatment without extracellular Ca^2+^ removal (***Figure 7-figure supplement 1A***), suggesting full alteration of intracellular Ca^2+^ distribution at later time of LP treatment. Since JNK contributes to inflammasome through ASC phosphorylation (Hara et al., 2013), we studied relationship between JNK and ASC phosphorylation/oligomerization. Intriguingly, ASC phosphorylation and oligomerization occurred 21 h after LP treatment but not 30 min after LPS treatment despite similar JNK activation (***Figure 7B and C***), suggesting that JNK activation 21 h after LP treatment leads to ASC phosphorylation and oligomerization likely due to signal 2 such as lysosomal events. Indeed, JNK activation by LP treatment for 21 h was markedly suppressed by *Kcnn4* or *Trpm2* KO (***Figure 7-figure supplement 1B***), supporting that events such as lysosomal Ca^2+^ release and/or K^+^ efflux are necessary for delayed JNK activation. SP600125, a JNK inhibitor, suppressed ASC phosphorylation and oligomerization by LP (***Figure 7D; Figure 7-figure supplement 1C***), indicating that JNK activation is necessary but not sufficient for ASC phosphorylation/oligomerization, as shown by no ASC phosphorylation/oligomerization 0.5 h after LPS treatment despite strong JNK activation. Consistent with roles of JNK in inflammasome activation by LP, IL-1β release by LP was inhibited by JNK inhibitor (***Figure 7E***).

**Figure 7.**
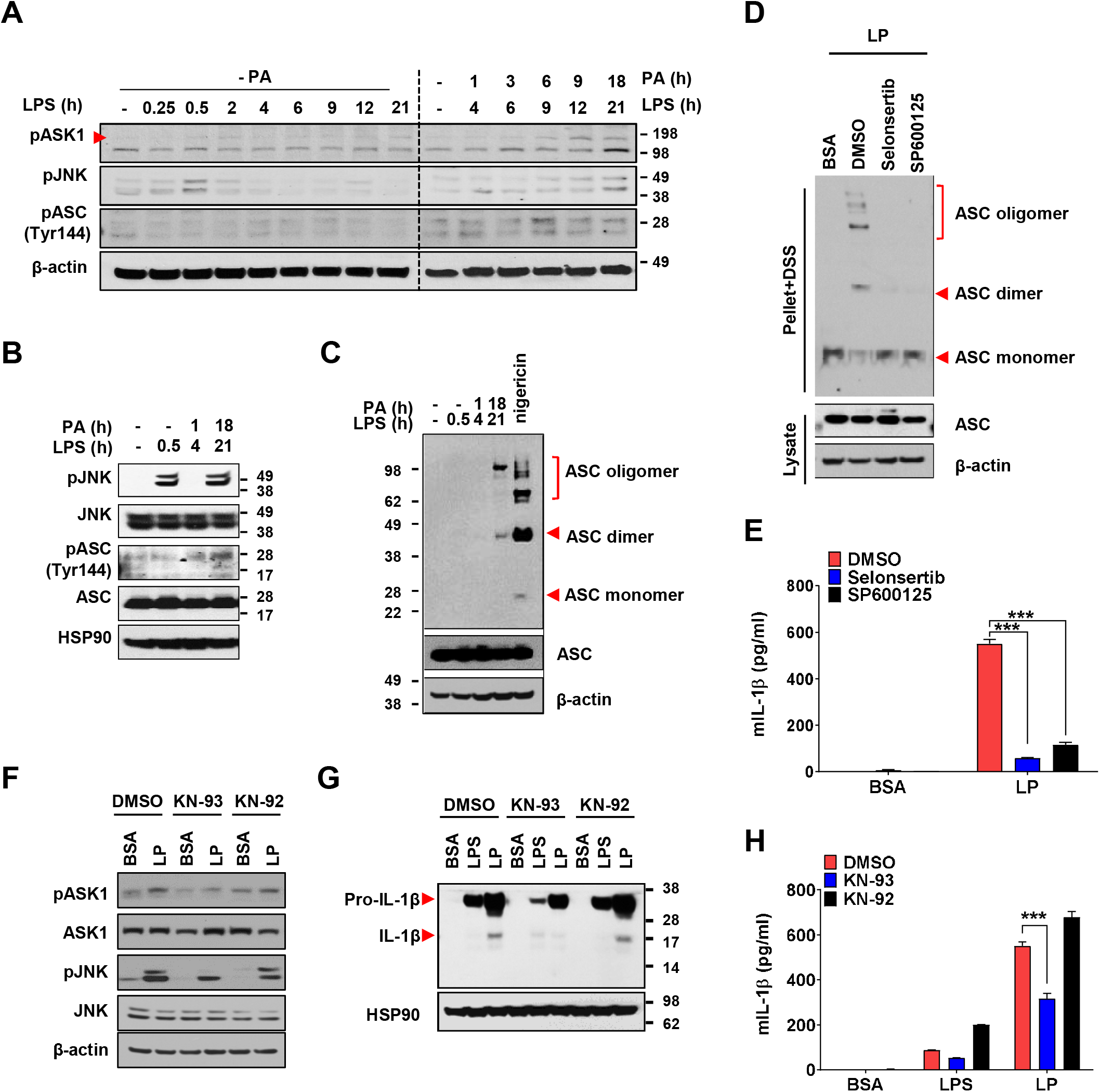
Mechanism of Ca^2+^-mediated inflammasome. (**A**) IB of BMDMs treated with LPS alone without PA (‘- PA’) for indicated time period (left half) or with ‘PA’ (together with LPS) for indicated time period after LPS pretreatment for 3 h (right half) (hence, the numbers indicating LPS treatment time in the right half is 3 + PA treatment time), using indicated Abs. (**B**) IB of BMDMs treated with LPS alone for 0.5 h or ‘PA’ (together with LPS) for 1 or 18 h after LPS pretreatment for 3 h (hence, the numbers indicating LPS treatment time of 4 or 21 h is 3 + PA treatment time), using indicated Abs. (**C**) BMDMs were treated with LPS alone for 0.5 h, ‘PA’ (together with LPS) for 1 or 18 h after LPS pretreatment for 3 h (hence, the numbers indicating LPS treatment time of 4 or 21 h is 3 + PA treatment time) or nigericin for 45 min after LPS pretreatment for 3 h. IB using indicated Abs after DSS crosslinking. (**D**) BMDMs were treated with LP for a total of 21 h including LPS pretreatment for 3 h in the presence or absence of ASK1 (selonsertib) or JNK inhibitor (SP600125). IB using indicated Abs after DSS crosslinking. (**E**) IL-1β ELISA of culture supernatant after treating BMDMs with LP for a total of 21 h including LPS pretreatment for 3 h in the presence or absence of selonsertib or SP600125. (**F-H**) IB using indicated Abs (F, G) and IL-1β ELISA of culture supernatant (H) after treating BMDMs with LPS alone for 21 h or with LP for a total of 21 h including LPS pretreatment for 3 h in the presence or absence of KN-93 or -92. (*n*=3). Data shown as means ± SEM from more than 3 independent experiments. ***p< 0.001 by two- way ANOVA with Tukey’s test (E, H). The online version of this article includes the following figure supplement for figure 7: **Figure supplement 1.** Effect of *Kcnn4* KO and inhibitor of TAK1 or ASK1 on activation of inflammasome and JNK.

When we studied roles of TAK1 as JNK upstream (Okada et al., 2014), IL-1β release by LP was not inhibited by 5Z-7-oxozeaenol, a TAK1 inhibitor (***Figure 7-figure supplement 1D***), in contrast to inflammasome activation by lysosomal rupture (Okada et al., 2014). We thus studied roles of ASK1 participating in inflammasome activation by diverse inflammasome activators such as bacteria or virus (Immanuel et al., 2019; Place et al., 2018). IL-1β release by LP was significantly inhibited by selonsertib, an ASK1 inhibitor (***Figure 7E***), suggesting roles of ASK1 in inflammasome activation by LP. Consistently, JNK activation by LP was inhibited by selonsertib (***Figure 7-figure supplement 1E***). Further, ASK1 activation was observed at later time of LP treatment (***Figure 7A***), suggesting that LP induces ASK1 activation likely through additional mechanisms such as lysosomal events at JNK upstream. Consistent with roles of ASK1 in JNK activation, ASK1 inhibitor blocked ASC phosphorylation/oligomerization as efficiently as JNK inhibitor (***Figure 7D; Figure 7*- *figure supplement 1C***), supporting that activated ASK1 induces JNK activation and subsequent ASC phosphorylation/oligomerization.

When we studied mechanism of ASK1 activation by LP, ASK1 activation by LP was attenuated by KN-93, a CaMKII inhibitor, but not by KN-92, a KN-93 congener without CaMKII inhibitory activity (***Figure 7F***), which supports roles of increased [Ca^2+^]_i_ and subsequent CAMKII in ASK1 activation by LP, similar to inhibition of lysosomal rupture-induced inflammasome by KN-93 (Okada et al., 2014). JNK phosphorylation and inflammasome activation as evidenced by maturation or release of IL-1β by LP were also inhibited by KN-93 but not by KN-92 (***Figure 7F-H***). These results indicate that increased [Ca^2+^]_i_ due to lysosomal Ca^2+^ release facilitated by ER→lysosome Ca^2+^ refilling and K^+^ efflux through KCa3.1 channel induces delayed activation of ASK1 and JNK, leading to ASC oligomerization or NLRP3 inflammasome by LP and metabolic inflammation.

## Discussion

Agents inducing lysosomal stress or lysosomotropic agents are well-known inflammasome activators (Hornung et al., 2008). Regarding mechanism of inflammasome by lysosomotropic agents, lysosomal Ca^2+^ has been incriminated (Okada et al., 2014; Weber and Schilling, 2014), while papers refuting the role of Ca^2+^ have been published (Katsnelson et al., 2015). We found that lysosomal Ca^2+^ release and subsequent [Ca^2+^]_i_ increase by LP lead to inflammasome through CAMKII-ASK1 and delayed JNK activation, which is critical for metabolic inflammation (***Figure 8***).

**Figure 8.**
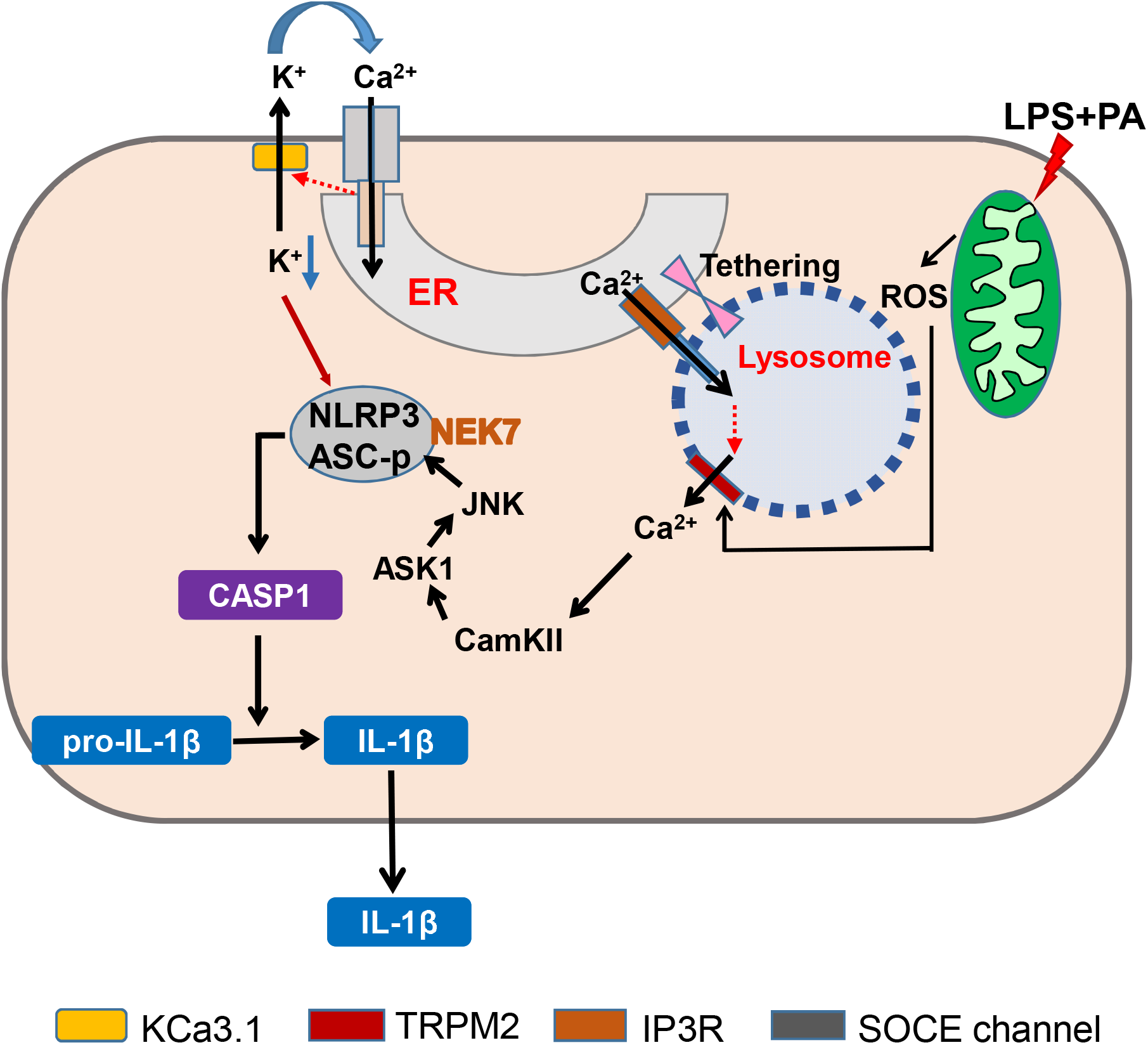
Graphic summary. LP, an effector combination activating inflammasome related to metabolic inflammation, induces generation of mitochondrial ROS, which activates TRPM2 channel on lysosome and releases lysosomal Ca^2+^. ER→lysosome Ca^2+^ refilling facilitated by ER-lysosome tethering replenishes diminished lysosomal Ca^2+^ content and supports sustained lysosomal Ca^2+^ release. ER emptying due to ER→lysosome Ca^2+^ refilling activates SOCE. SOCE, in turn, is positively modulated by K^+^ efflux through KCa3.1, a Ca^2+^-activated K^+^ efflux channel, mediating hyperpolarization-induced acceleration of extracellular Ca^2+^ influx. Ca^2+^ release from lysosome activates CaMKII, which induces delayed activation of ASK1 and JNK. Delayed JNK activation leads to ASC phosphorylation and oligomerization, leading to formation of inflammasome complex together with NLRP3 and NEK7. K^+^ efflux changes intracellular milieu and induces structural changes of NLRP3 or NLRP3 binding to PI(4)P on dispersed Golgi network, facilitating inflammasome activation. Golgi complex and microtubule-organizing center (MTOC) are not shown for clarity. (CASP1, caspase 1)

Among diverse inflammasome activators, LP is a representative effector combination responsible for metabolic inflammation related to metabolic syndrome. However, mechanism of inflammasome activation by LP has been unclear. It is well known that PA induces stresses of organelle such as ER and mitochondria (Bachar et al., 2009; Korge et al., 2003). We observed that mitochondrial ROS accumulate by LP and activate lysosomal Ca^2+^ release through TRPM2. Roles of mitochondrial ROS in inflammasome activation has been reported (Weber and Schilling, 2014), however, causal relationship between mitochondrial ROS and lysosomal events has not been addressed. Roles of TRPM2 in diabetes or β-cell function have been studied (Uchida and Tominaga, 2011; Zhang et al., 2012). However, roles of TRPM2 in metabolic inflammation was not studied.

While we have shown roles of TRPM2 in inflammasome and metabolic inflammation, TRPM2 exists on both plasma membrane and lysosome (Wang et al., 2020). Roles of plasma membrane TRPM2 in inflammasome by high glucose or particulate materials have been reported (Tseng et al., 2017; Zhong et al., 2013), while the role of TMRP2 in inflammation activation by lipid stimulators related to metabolic inflammation has not been addressed. However, we observed no activation of TRPM2 current on the plasma membrane of Mφs treated with LP, arguing against roles of plasma membrane TRPM2 in inflammasome by LP.

Further, dissipation of lysosomal Ca^2+^ reservoir by bafilomycin A1 abrogated [Ca^2+^]_i_ increase by LP, strongly supporting roles of lysosomal TRPM2 rather than plasma membrane TRPM2 in inflammation by LP.

We observed decreased [Ca^2+^]_ER_ in inflammasome activation by LP, which was due to ER→lysosome Ca^2+^ refilling through IP_3_R to replenish diminished [Ca^2+^]_Lys_ after lysosomal Ca^2+^ release. Previous papers have shown roles of ER Ca^2+^ in inflammasome activation (Lee et al., 2012), however, role of ER Ca^2+^ replenishing reduced [Ca^2+^]_Lys_ in inflammasome has not been addressed. Direct Ca^2+^ efflux from ER to cytoplasm which was reported in inflammasome activation by ATP (Lee et al., 2012) cannot explain ER→lysosome refilling or abrogation of LP-induced increase of [Ca^2+^]_i_ by bafilomycin A1 which would not decrease but increase [Ca^2+^]_i_ after ER Ca^2+^ release into cytoplasm due to abrogated lysosomal Ca^2+^ buffering (Sanurjo et al., 2014). Likely due to ER Ca^2+^ release to replenish reduced [Ca^2+^]_Lys_, [Ca^2+^]_ER_ was reduced, which in turn induced SOCE. Then, extracellular Ca^2+^ influx appears to replenish ER Ca^2+^ after ER Ca^2+^ loss due to ER→lysosome Ca^2+^ refilling. Roles of extracellular Ca^2+^ influx in inflammasome have been suggested, while extracellular Ca^2+^ influx was unrelated to ER Ca^2+^ store (Lee et al., 2012; Tseng et al., 2017). STIM1 aggregation and its colocalization with ORAI1 strongly support SOCE in inflammasome by LP to replenish ER Ca^2+^ depletion (Vaca, 2010), which is different from activation of Ca^2+^- sensing receptor by extracellular Ca^2+^ reported in ATP-induced inflammasome (Lee et al., 2012). We have observed that BTP2 inhibiting Ca^2+^ influx through ORAI1 (Bogeski et al., 2010) suppressed LP-induced inflammasome, suggesting that extracellular Ca^2+^ influx is a process of SOCE through STIM1/ORAI1 channel activated by ER Ca^2+^ depletion rather than a direct extracellular Ca^2+^ influx into cytoplasm through TRPM2 on the plasma membrane (Zhong et al., 2013). Thus, these results additionally support roles of lysosomal TRPM2 but not plasma membrane TRPM2 in LP-induced inflammasome.

We observed that K^+^ efflux occurs in inflammasome by LP, similar to that by other activators. K^+^ efflux through KCa3.1 channel appears to induce NLRP3 binding to NEK7 and oligomerization which was inhibited by TRAM-34 or *Kcnn4* KO. Previous papers suggested roles of KCa3.1 channel in SOCE, while such relationship was unrelated to inflammasome (Duffy et al., 2015; Gao et al., 2010). A KCa3.1 channel activator together with LPS has been reported to induce IL-1β release; however, roles of KCa3.1 channel in authentic inflammasome have not been shown (Schroeder et al., 2017). Although we observed significant roles of KCa3.1 channel in inflammasome activation, contribution of other Ca^2+^- activated K^+^ channels such as BK channels cannot be eliminated since iberitoxin, a BK channel inhibitor, has been reported to inhibit ATP-induced inflammasome (Schroeder et al., 2017). However, relationship between BK channel and Ca^2+^ flux was not studied. Previous papers also reported roles of TWIK2 channel, a two-pore K^+^ channel, in K^+^ efflux in ATP-induced inflammasome, which was inhibited by quinine (Di et al., 2017). Role of THIK-1, another two-pore K^+^ channel in IL-1β release from microglia by ATP has also been reported (Madry et al., 2018). However, inflammasome by LP was not inhibited by quinine. Further, relationship between K^+^ efflux through two-pore K^+^ channels and SOCE was not studied. Relationship between ATP-induced P2X7 activation and K^+^ efflux could be distinct from coupling of Ca^2+^ influx and K^+^ efflux by lysosomotropic agents or other events primarily affecting lysosomal Ca^2+^ channel (Di et al., 2017; Muñoz-Planillo et al., 2013). Such differences could explain no effect of quinine on LP-induced inflammasome or undiminished lysosomotropic agents-induced inflammasome in *THIK-1*-KO Mφs or microglia (Drinkall et al., 2022).

Altogether, we have shown the sequential events from mitochondrial ROS-induced lysosomal TRPM2 channel activation to ER→lysosome Ca^2+^ refilling and SOCE coupled with KCa3.1 K^+^ efflux channel activation in inflammasome by LP and metabolic inflammation (***Figure 8***), which could also be applied to that by other lysosomotropic agents. These results suggest mitochondrial ROS-lysosomal TRPM2 axis as a potential therapeutic target for treatment of metabolic inflammation. Elucidation of KCa3.1 channel as the K^+^ efflux channel in inflammasome and its role in facilitation of extracellular Ca^2+^ influx through SOCE would provide another target for modulation of inflammasome that is activated in a variety of diseases or conditions in addition to metabolic syndrome. ER→lysosome refilling might have implication in diverse conditions or diseases associated with lysosomal Ca^2+^ changes such as autophagy, vaccination or inflammation (Park et al., 2022; Tahtinen et al., 2022).

## Materials and Methods

### GCaMP3 Ca^2+^ imaging

Bone marrow-derived Mφs (BMDMs) (kindly provided by Je-Wook Yu, Yonsei University) grown on 4-well chamber were transfected with a plasmid encoding a perilysosomal GCaMP3-ML1 Ca^2+^ probe (Shen et al., 2012). After 48 h, cells were treated with LP for 1 h after LPS pretreatment for 3 h, and then lysosomal Ca^2+^ release was measured in a basal Ca^2+^ solution containing 145 mM NaCl, 5 mM KCl, 3 mM MgCl_2_, 10 mM glucose, 1 mM EGTA and 20 mM HEPES (pH 7.4) by monitoring fluorescence intensity at 470 nm using a LSM780 confocal microscope (Carl Zeiss, LSM 780). GPN was added at the indicated time points.

### Determination of [Ca^2+^]_i_

After pretreatment with LPS for 3 h and treatment with LP for 1 h, cells were loaded with Fluo-3-AM (Invitrogen) at 37°C for 30 min. [Ca^2+^]_i_ was measured in a basal Ca^2+^ solution containing 145 mM NaCl, 5 mM KCl, 3 mM MgCl_2_, 10 mM glucose, 1 mM EGTA and 20 mM HEPES (pH 7.4), using a LSM780 confocal microscope (Carl Zeiss).

For ratiometric determination of [Ca^2+^]_i_, cells treated with LP were loaded with 2 μM of the acetoxymethyl ester form of Fura-2 (Invitrogen) in RPMI-1640 at 37°C for 30 min and fluorescence data were analyzed using MetaFluor (Molecular Devices) on Axio Observer A1 (Zeiss) equipped with 150 W xenon lamp Polychrome V (Till Photonics), CoolSNAP-Hq2 digital camera (Photometrics), and Fura-2 filter set. Fluorescence at 340/380 nm was measured in phenol red-free RPMI, and converted to [Ca^2+^]_i_ using and the following equation (Grynkiewicz et al., 1985).

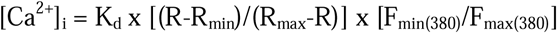

where K_d_ = Fura-2 dissociation constant (224 nM at 37°C), F_min(380)_ = 380 nm fluorescence in the absence of Ca^2+^, F_max(380)_ = 380 nm fluorescence with saturating Ca^2+^, R = 340/380 nm fluorescence ratio, R_max_ = 340/380 nm ratio with saturating Ca^2+^, and R_min_ = 340/ 380 nm ratio in the absence of Ca^2+^.

### Measurement of [Ca^2+^]_Lys_

To measure [Ca^2+^]_Lys_, cells were loaded with 100 μg/ml OGBD, an indicator of lysosomal luminal Ca^2+^, at 37°C in the culture medium for 12 h to allow uptake by endocytosis. After additional incubation for 4 h without indicator, cells were treated with LP for 1 h after LPS pretreatment for 3 h, and then washed in HBS (135 mM NaCl, 5.9 mM KCl, 1.2 mM MgCl_2_, 1.5 mM CaCl_2_, 11.5 mM glucose, 11.6 mM HEPES, pH 7.3) for confocal microscopy (Garrity et al., 2016).

### Measurement of ER Ca^2+^ content ([Ca^2+^]_ER_)

BMDMs grown on 4-well chamber were transfected with *GEM-CEPIA1er* (Addgene) (Suzuki et al., 2014) or a ratiometric FRET-based Cameleon probe *D1ER* (Park et al., 2009) using Lipofectamine 2000. After 48 h, cells were pretreated with LPS for 3 h and then treated with LP for 1 h in a Ca^2+^-free Krebs-Ringer bicarbonate (KRB) buffer (Sigma) to eliminate the effect of extracellular Ca^2+^ influx into ER (Xu et al., 2015). Fluorescence was measured using an LSM780 confocal microscope (Zeiss) at an excitation wavelength of 405 nm and an emission wavelength of 466 or 520 nm. F466/F520 was calculated as an index of [Ca^2+^]_ER_ (Suzuki et al., 2014). D1ER fluorescence intensity ratio (F540/F490) was determined using an LSM780 confocal microscope (Zeiss).

### Simultaneous monitoring of [Ca^2+^]_Lys_ and [Ca^2+^]_ER_

Twenty-four h after transfection with *GEM-CEPIA1er*, BMDMs were loaded with OGBD for 12 h and chased for 4 h. After treating cells with LP for a total of 4 h including LPS pretreatment for 3 h, medium was changed to a fresh one without LP. Cells were then monitored for [Ca^2+^]_Lys_ and [Ca^2+^]_ER_ in a Ca^2+^-free KRB buffer using an LSM780 confocal microscope.

### Proximity ligation assay (PLA)

Contact between ER and lysosome was examined using Duolink In Situ Detection Reagents Red kit (Sigma) according to the manufacturer’s protocol. Briefly, BMDMs treated with test agents were incubated with antibodies (Abs) to ORP1L (Abcam, 1:200) and VAP-A (Santa Cruz, 1:200), or with those to KCNN4 (Invitrogen) and ORAI1 (Novusbio) at 4°C overnight. After washing, cells were incubated with PLA plus and minus probes at 37°C for 1 h. After ligation reaction to close the circle and rolling circle amplification (RCA) of the ligation product, fluorescence-labelled oligonucleotide hybridized to RCT product was observed by fluorescence microscopy.

### SOCE channel activation

BMDMs transfected with *YFP-STIM1* and *3xFLAG-mCherry Red-Orai1/P3XFLAG7.1* (kindly provided by Joseph Yuan, University of North Texas Health Science Center, USA through Cha S-G, Yonsei University) were treated with LP for a total of 4 h including LPS pretreatment for 3 h, which were then subjected to confocal microscopy to visualize STIM1 puncta and their co-localization with ORAI1.

### Abs and immunoblotting (IB)

Cells or tissues were solubilized in a lysis buffer containing protease inhibitors. Protein concentration was determined using Bradford method. Samples (10∼30 μg) were separated on 4∼12% Bis-Tris gel (NUPAGE, Invitrogen), and transferred to nitrocellulose membranes for IB using the ECL method (Dongin LS). For IB, Abs against the following proteins were used: IL-1β (R&D systems, AF-401-NA, 1: 1,000), caspase 1 p20 (Millipore, ABE1971, 1: 1,000), ASC (Adipogen AL177, 1: 1,000), phospho-JNK (Cell signaling #9251, 1: 1,000), JNK (Santa Cruz sc7345, 1: 1000), phospho-ASK1 (Invitrogen PA5-64541, 1: 1,000), ASK1 (Abcam ab45178, 1: 1,000), phospho-ASC (ECM Biosciences AP5631, 1: 1,000), NLRP3 (Invitrogen MA5-23919, 1:1000), NEK7 (Abcam ab133514, 1:1000), HSP 90 (Santa Cruz sc13119, 1: 1,000) and β-actin (Santa Cruz sc47778, 1: 1,000).

### Immunoprecipitation

After lysis of cells in an ice-cold lysis buffer (400[mM NaCl, 25[mM Tris-HCl, pH 7.4, 1[mM EDTA, and 1% Triton X-100) containing protease and phosphatase inhibitors, lysates were centrifuged at 12,000[*g* for 10 min in microfuge tubes, and supernatant was incubated with anti-NEK7 (Abcam, 1:1000) Ab or control IgG in binding buffer (200[mM NaCl, 25[mM Tris-HCl, pH 7.4, 1[mM EDTA) with constant rotation at 4°C for 1 h. After adding 50 μl of 50% of Protein-G bead (Roche Applied Science) to lysates and incubation with rotation at 4°C overnight, resins were washed with binding buffer. After resuspending pellet in a sample buffer (Life Technology) and heating at 100°C for 3 min, supernatant was collected by centrifugation at 12,000[*g* for 30[sec, followed by electrophoretic separation in a NuPAGE^®^ gradient gel (Life Technology). IB was conducted by sequential incubation with anti-NEK7 or -NLRP3 Ab as the primary Ab and horseradish peroxidase-conjugated anti-rabbit IgG or -mouse IgG. Bands were visualized using an ECL kit.

### Detection of ASC oligomerization

BMDMs were washed in ice-cold PBS, and then lysed in NP-40 buffer (20 mM HEPES-KOH pH 7.5, 150 mM KCl, 1% NP-40, and protease inhibitors). Lysate was centrifuged at 2,000 *g*, 4°C for 10 min. Pellets were washed and resuspended in PBS containing 2 mM disuccinimidyl suberate (DSS) for crosslinking, followed by incubation at room temperature for 30 min. Samples were then centrifuged at 2,000 *g*, 4°C for 10 min. Precipitated pellets and soluble lysates were subjected to IB using anti-ASC Ab.

### Blue Native PAGE

Blue Native polyacrylamide gel electrophoresis (BN-PAGE) was performed using Bis-Tris NativePAGE system (Invitrogen, Carlsbad, CA, USA), according to the manufacturer’s instructions. Briefly, Cells were collected and lysed in 1 x NativePAGE Sample Buffer (Invitrogen) containing 1% digitonin and protease inhibitor cocktail, followed by centrifugation at 13,000 rpm, 4°C for 20 min. 20 μl supernatant mixed with 1 μl 5% G-250 Sample Additive was loaded on a NativePAGE 3∼12% Bis-Tris gel. Samples separated on gels were transferred to PVDF membranes (Millipore, Darmstadt, Germany) using transfer buffer, followed by IB using anti-NLRP3 Ab.

### Immunofluorescence study

Cells were grown on 4-chamber plates. After treatments, cells were fixed with 4% paraformaldehyde for 15 min and permeabilized with 0.5% triton X-100 for 15 min. After blocking with 5% goat serum for 1 h, cells were incubated with anti-ASC at 4°C overnight. On the next day, samples were incubated with Alexa 488-conjugated anti-mouse or anti-rabbit IgG Ab (Invitrogen) for 1 h. After nuclear staining with DAPI (Invitrogen), cells were subjected to confocal microscopy (Carl Zeiss, LSM 780).

### Measurement of [K^+^]_i_

BMDMs treated with LP for a total of 21 h including LPS pretreatment for 3 h, were labeled with 5 μM Asante Potassium Green-2-AM (Abcam) at 37 °C for 30 min. After washing twice with PBS, cells were subjected to confocal microscopy (Carl Zeiss, LSM 780).

### ELISA of cytokines

Cytokine content in culture supernatants of BMDMs or peritoneal Mφs was determined using mouse ELISA kits (R&D Systems), according to the manufacturer’s instruction.

### RNA extraction and real-time RT-PCR

Total RNA was extracted from cells or tissues using TRIzol (Invitrogen), and cDNA was synthesized using MMLV Reverse Transcriptase (Promega), according to the manufacturer’s protocol. Real-time RT-PCR was performed using SYBR green (Takara) in QuantStudio3 Real-Time PCR System (Applied Biosystems). All expression values were normalized to *GAPDH* or *Rpl32* mRNA level.

Primer sequences are as follows: *Kcnn4*-F, 5’-AACTGGCATCGGACTCATGGTT-3’; *Kcnn4*-R, 5’-AGTCATGAACAGCTGGACCTC-3’.

### Animals

Eight-weeks-old male *Trpm2-*KO mice (kindly provided by Yasuo Mori, Kyoto University, Japan) were maintained in a 12-h light/12-h dark cycle and fed HFD for 12 weeks. During the observation period, mice were monitored for glucose profile and weighed. *Kcnn4*-KO mice were from Jackson Laboratories. All animal experiments were conducted in accordance with the Public Health Service Policy in Humane Care and Use of Laboratory Animals. Mouse experiments were approved by the IACUC of the Department of Laboratory Animal Resources of Yonsei University College of Medicine, an AAALAC-accredited unit.

### Cell culture and drug treatment

BMDMs were cultured in DMEM supplemented with 10% fetal bovine serum (FBS), 100 U/ml penicillin and 100 μg/ml streptomycin (Lonza). For drug treatment, the following concentrations were used: 50 or 100 ng/ ml LPS (Sigma), 300 μM PA (Sigma), 50 μM BAPTA-AM (Invitrogen), 10 μM nigericin (Sigma), 3 mM ATP (Roche), 25 μg/ml MSU (InvivoGen), 0.3 mM LLOMe (Sigma), 3 μM Xestospongin C (Abcam), 10 μM dantrolene (Sigma), 10 μM TPEN (Sigma), 3 mM EGTA (Sigma), 100 nM 2-APB (Sigma), 10 μM BTP2 (Merk Millipore), 5 mM NAC (Sigma), 100 μM MitoTEMPOL (Sigma), 10 μM selonsertib (Selleckchem), 10 μM SP600125 (Sigma), 100∼500 nM 5Z-7-oxozeaenol (Sigma), 10 μM KN-93 (Tocris), 10 μM KN-92 (Tocris), 30 μM apigenin (Sigma), quercetin (Sigma), 1 μM bafilomycin A1, 10 μM TRAM-34 (Sigma), 10 μM ACA (Sigma), 100 nM apamin (Sigma), 1 μM paxillin (Sigma), 100 μM quinine (Sigma), 1 mM barium sulfate (Sigma), 10 nM UCL 1684 (Sigma), and 10 μM 4-AP (Sigma). PA stock solution (50 mM) was prepared by dissolving in 70% ethanol and heating at 55°C. Working solution was made by diluting PA stock solution in 2% fatty acid-free BSA-DMEM or -RPMI.

### Peritoneal Mφs

Mice were injected intraperitoneally with 3 ml of 3.85% Brewer thioglycollate. Three days after injection, peritoneal Mφs were harvested, seeded at 6-well plates and maintained in RPMI containing 10% FBS, 100 U/ml penicillin and 100 μg/ml streptomycin at 37°C for 24 h in a humid atmosphere of 5% CO_2_ before treatment. Peritoneal Mφs were used as the primary Mφs throughout the study.

### Stromal vascular fraction (SVF)

SVF of epididymal adipose tissue was prepared as described (Lee et al., 2016). Briefly, after cutting SVF into small pieces and incubation in a 2 mg/ml collagenase D solution (Roche) at 37°C in a water bath for 45 min, digested tissue was centrifuged at 1,000 *g* for 8 min. After filtering through a 70-μm mesh and lysis of RBC, SVF was suspended in PBS supplemented with 2% BSA (Sigma), 2 mM EDTA (Cellgro) for further experiments.

### Metabolic studies

IPGTT was performed by intraperitoneal injection of 1 g/kg glucose solution after overnight fasting. Blood glucose concentrations were determined using One Touch glucometer (LifeScan) before (0 min) and 15, 30, 60, 120 and 180 min after glucose injection. Serum insulin was measured using Mouse Insulin ELISA kit (TMB) (AKRIN-011T, Shibayagi, Gunma, Japan). HOMA-IR index was calculated according to the following formula: [fasting insulin (μlU/ml) × fasting glucose (mg/dl)]/405.

### Histology and immunohistochemistry

Tissue samples were fixed with 10% buffered formalin and embedded in paraffin. Sections of 5 μm thickness were stained with H&E for morphometry, or immunostained with F4/80 Ab (Abcam) to detect Mφ aggregates surrounding adipocytes (crown-like structures, CLS).

### Intracellular ROS

To determine ROS, cells were treated with LPS alone for 21 h, PA alone for 21 h or LP for a total of 21 h including LPS pretreatment for 3 h in the presence or absence of 5 mM NAC. After incubation with 5 μM CM-H2DCFDA (Invitrogen) at 37°C for 30 min in culture media without FBS and recovery in a complete media for 10 min, confocal microscopy was conducted. To study mitochondria-specific ROS, cells were treated with LPS alone for 21 h, PA alone for 21 h or LP for a total of 21 h including LPS pretreatment for 3 h in the presence or absence of 100 μM MitoTEMPOL. After staining with 5 μM MitoSOX (Invitrogen) at 37°C for 30 min and suspension in PBS-1% FBS, cells were subjected to flow cytometry on FACSVerse (BD Biosciences), and data were analyzed using FlowJo software (TreeStar).

### Measurement of KCa3.1 and TRPM2 current

To measure KCa3.1 channel activity with intact cytosolic Ca^2+^ environment, nystatin perforated whole-cell patch was performed. 180∼250 μg/ml of nystatin was added to intracellular pipette solution (140 mM KCl, 5 mM NaCl, 0.5 mM MgCl_2_, 3 mM MgATP, and 10 mM HEPES, pH adjusted to 7.3 with KOH). The extracellular bath solution (145 mM NaCl, 3.6 mM KCl, 1.3 mM CaCl_2_, 1 mM MgCl_2_, 5 mM glucose, 10 mM HEPES, and 10 mM sucrose, pH adjusted to 7.3 with NaOH) was perfused through recording chamber. After perforated whole-cell configuration is formed, KCa3.1 current was activated by applying ramp-pulse voltage from -120 to 60 mV. Voltage-dependent K^+^ channels were suppressed by - 10 mV holding potential. Intrinsic KCa3.1 current activity was isolated by application of a selective inhibitor, TRAM-34. Activity of TRAM-34-sensitive current was analyzed with slope conductance fitting between -60 to -40 mV.

To measure plasma TRPM2 channel activity, conventional whole-cell patch was performed. To induce TRPM2 current, 200 μM of ADP-ribose was added to intracellular pipette solution (87 mM Cs-glutamate, 38 mM CsCl, 10 mM NaCl, 10 mM HEPES, 1 mM EGTA, and 0.9 mM CaCl_2_, pH adjusted to 7.2 with CsOH). Cells were perfused with extracellular bath solution (143 mM NaCl, 5.4 mM KCl, 0.5 mM MgCl_2_, 1.8 mM CaCl_2_, 10 mM HEPES, 0.5 mM NaH_2_PO_4_, and 10 mM glucose, pH 7.4 adjusted with NaOH) during TRPM2 current recording. When maximal TRPM2 current was established, 20 μM ACA was added to inhibit TRPM2 current.

Patch clamp experiments were performed at room temperature. Patch clamp pipettes were pulled to resistances of 2∼3 MΩ with PP-830 puller (Narishige, Japan). Electrophysiological data was recorded and analyzed with Axopatch 200B, Digidata 1440A, and Clampfit 11 program (Axon Instruments, CA).

### Statistical analysis

All values are expressed as the means ± s.e.m. from ≥ 3 independent experiments performed in triplicate. Statistical significance was tested with two-tailed Student’s *t* test to compare values between two groups. One-way ANOVA with Tukey’s test was employed to compare between multiple groups. Two-way ANOVA with Bonferroni, Sidak or Turkey’s test were employed to compare multiple repeated measurements between groups. All analyses were performed using GraphPad Prism Version 8 software (La Jolla, CA, USA). p values less than 0.05 were considered to represent statistically significant differences.

## Acknowledgements

This study was supported by the Basic Science Research Program through the National Research Foundation of Korea funded by the Ministry of Education (NRF- 2019R1I1A1A01063850 to H.K.), and National Research Foundation of Korea (NRF) grant funded by the Korea government (MSIT) (NRF-2019R1A2C3002924 and Rc-2023- 00219563 to M.-S.L.). M-S Lee is the recipient of Korea Drug Development Fund from the Ministry of Science & ICT, Ministry of Trade, Industry & Energy and Ministry of Health & Welfare (HN22C0278), and a grant from the Bio&Medical Technology Development Program (2017M3A9G7073521).

## Author contributions

M.-S. L. conceived the experiment. H. K., S.W.C., J.Y.K., S-J.O. and S.J.K. conducted experiment. M.-S. L., H. K., S.W.C. and S.J. K. wrote the manuscript.

---

**Figure 1−figure supplement 1.**
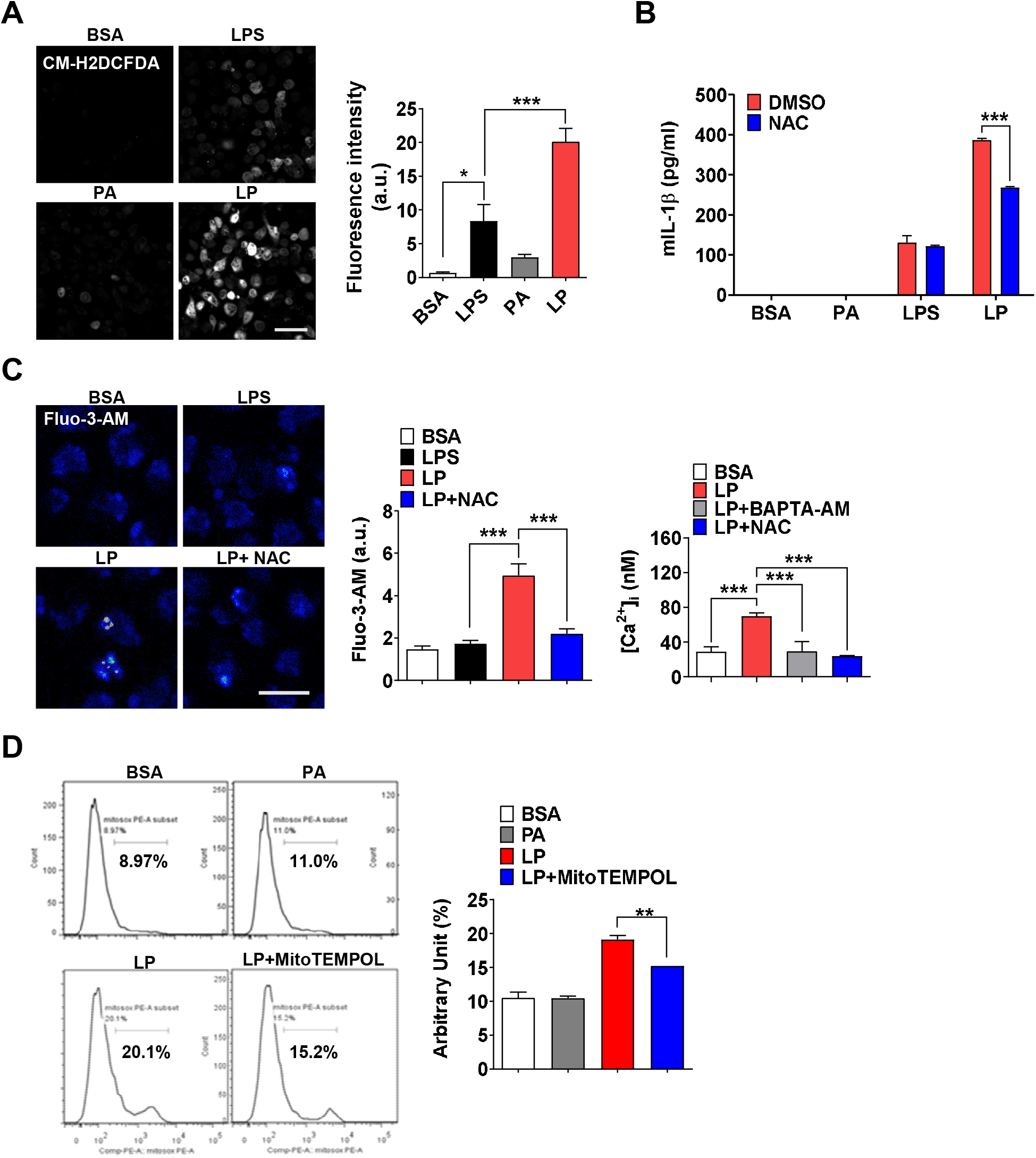
Mitochondrial ROS in inflammasome activation by LPS+PA (LP). (**A**) Cellular ROS in peritoneal Mφs treated with LPS alone or PA alone for 21 h, or LP for a total of 21 h including LPS pretreatment for 3 h, determined by confocal microscopy after CM-H2DCFDA loading (right). Representative confocal images (left) (scale bar, 50 μm). (**B**) IL-1β ELISA of culture supernatant after treating Mφs with LPS alone or PA alone for 21 h, or with LP for a total of 21 h including LPS pretreatment for 3 h in the presence or absence of NAC (*n*=3). (**C**) [Ca^2+^]_i_ in Mφs treated with LPS alone for 4 h or with LP for a total of 4 h including LPS pretreatment for 3 h in the presence or absence of NAC, determined by confocal microscopy after Fluo-3-AM loading (middle) or ratiometric measurement after Fura-2 loading (right). Representative Fluo-3 fluorescence images (left) (scale bar, 20 μm) (*n*=13 for BSA, Fluo-3-AM; *n*=12 for LPS, Fluo-3-AM; *n*=26 for LP, Fluo-3-AM; *n*=17 for LP+NAC, Fluo-3-AM; *n*=19 for BSA, Fura-2; *n*=23 for LP, Fura-2; *n*=23 for LP+BAPTA-AM, Fura-2; *n*=21 for LP+NAC, Fura-2). (**D**) Mitochondrial ROS in Mφs treated with PA alone for 21 h or with LP for a total of 21 h including LPS pretreatment for 3 h in the presence or absence of MitoTEMPOL, determined by flow cytometry after MitoSOX staining (right). Representative histograms (left) (*n*=3). Data shown as means ± SEM from more than 3 independent experiments. *p < 0.05, **p < 0.01 and ***p < 0.001 by one-way ANOVA with Tukey’s test (A, C, D), or two-way ANOVA with Sidak test (B).

**Figure 2−figure supplement 1.**
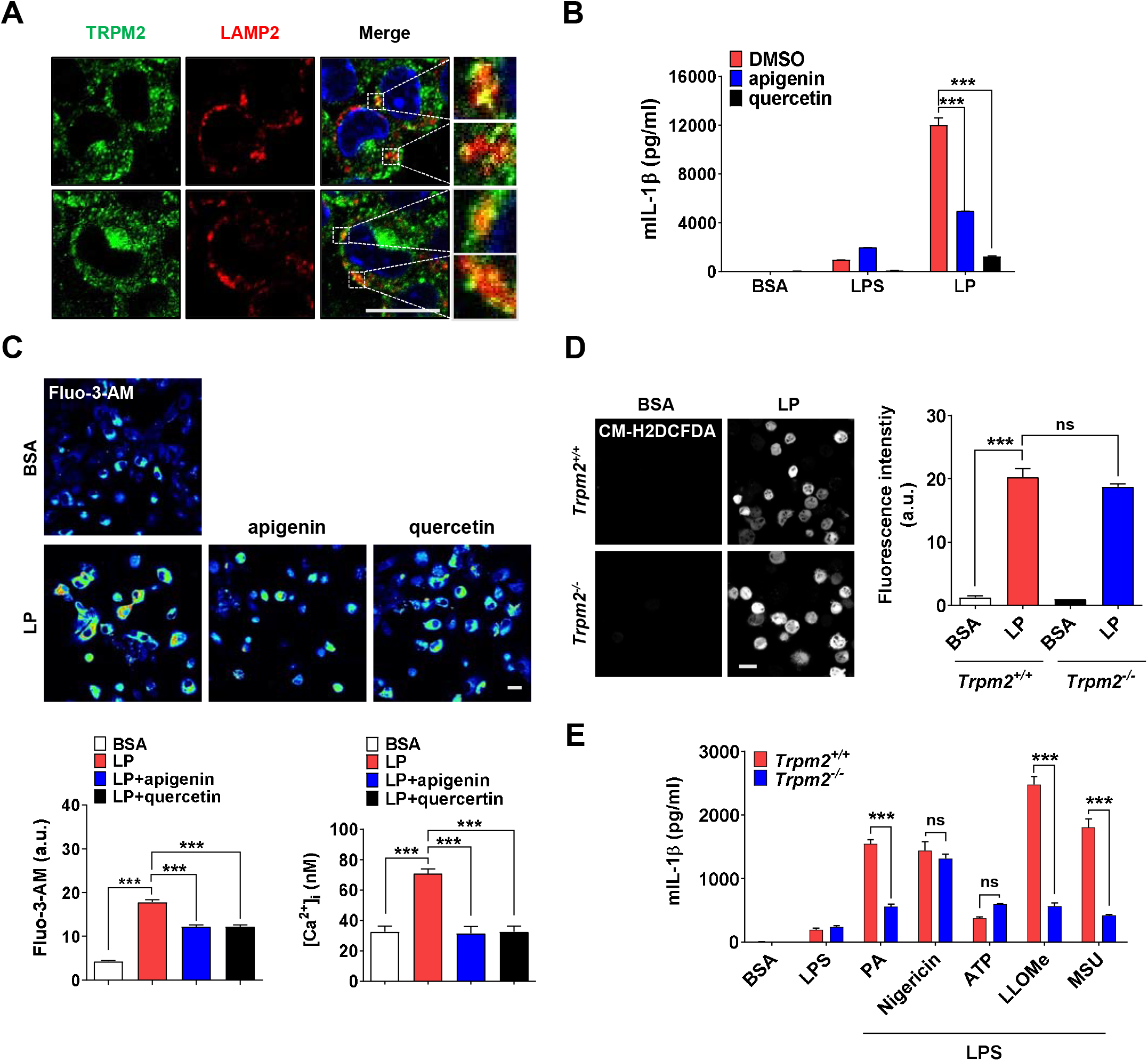
Effect of CD38 inhibitors and *Trpm2* KO on inflammasome. (**A**) Confocal microscopy was conducted after immunofluorescence staining of BMDMs using anti-TRPM2 and LAMP2 Abs. Yellow spots indicate TRPM2 on lysosome. (Rectangles were magnified) (**B**) IL-1β ELISA of culture supernatant after treating Mφs with LPS alone for 21 h or with LP for a total of 21 h including LPS pretreatment for 3 h in the presence or absence of apigenin or quercetin (*n*=3). (**C**) [Ca^2+^]_i_ in Mφs treated with LPS alone for 4 h or with LP for a total of a 4 h including LPS pretreatment for 3 h in the presence or absence of apigenin or quercetin, determined by confocal microscopy after Fluo-3-AM loading (lower left) or ratiometric measurement after Fura-2 loading (lower right). Representative Fluo-3 fluorescence images (upper) (*n*=4 for BSA, Fluo-3-AM; *n*=4 for LP, Fluo-3-AM; *n*=4 for LP+apigenin, Fluo-3-AM; *n*=4 for LP+quercetin, Fluo-3-AM; *n*=13 for BSA, Fura-2; n=16 for LP, Fura-2; *n*=20 for LP+apigenin, Fura-2; *n*=13 for LP+quercetin, Fura-2). (**D**) Cellular ROS in Mφs from Trpm2^+/+^ or Trpm2^-/-^ mice treated with LP for a total of 21 h including LPS pretreatment for 3 h, determined by CM-H2DCFDA staining (right). Representative fluorescence images (left) (*n*=4). (**E**) IL-1β ELISA of culture supernatant of Mφs from Trpm2^+/+^ or Trpm2^-/-^ mice treated with PA, nigericin, ATP, L-leucyl-L-leucine methyl ester (LLOMe) and MSU for 18 h, 45 min, 1 h, 45 min and 3 h, respectively, after LPS pretreatment for 3 h (*n*=3). Data shown as means ± SEM from more than 3 independent experiments. ***p < 0.001 by one-way ANOVA with Tukey’s test (C,D) or two-way ANOVA with Tukey’s test (B,E). Scale bar, 20 μm.

**Figure 3−figure supplement 1.**
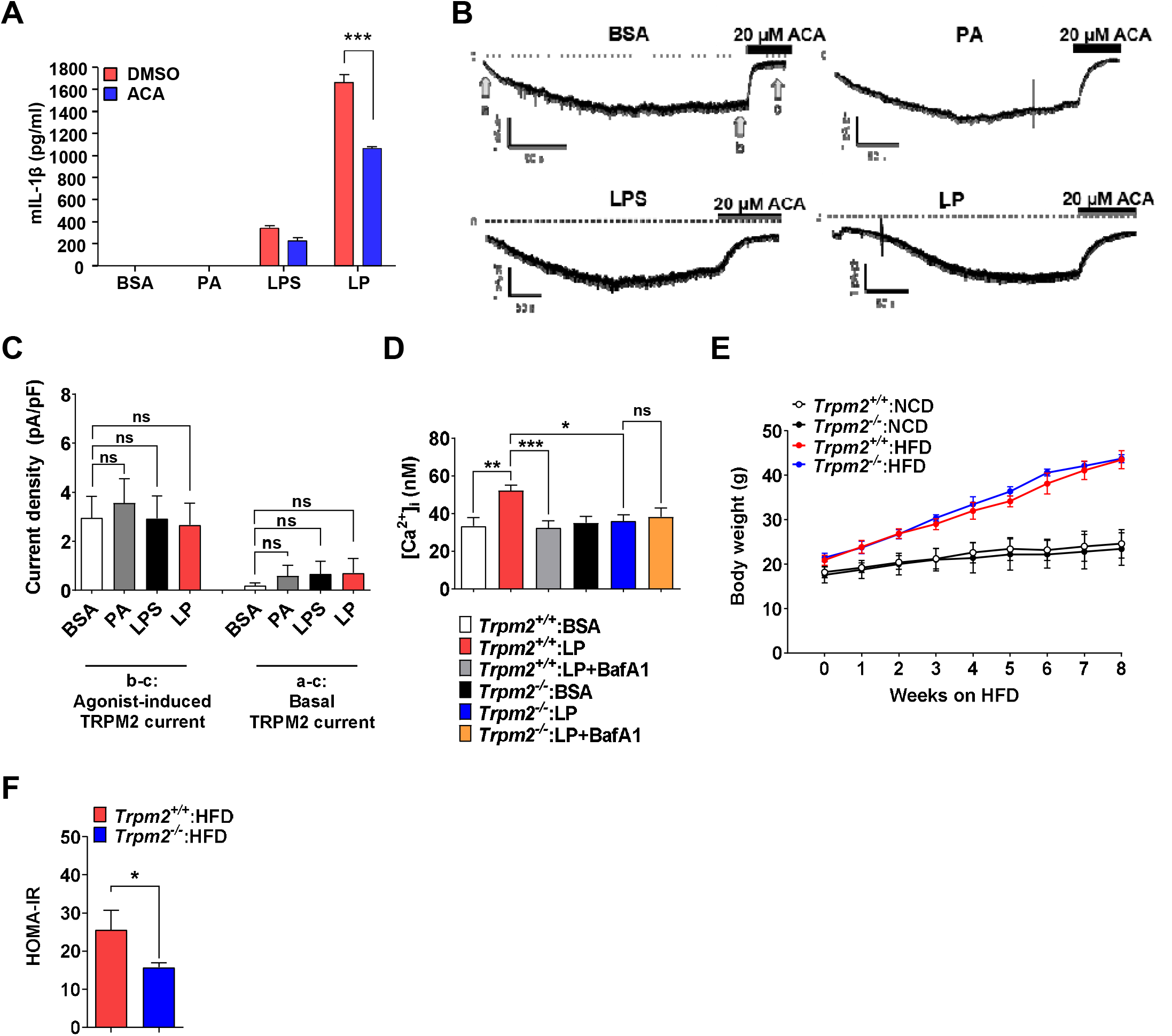
Plasma membrane TRPM2 current in Mφs and metabolic profile of *Trpm2*-KO mice. (**A**) IL-1β ELISA of culture supernatant after treating Mφs with LPS alone or PA alone for 21 h, or with LP for a total of 21 h including LPS pretreatment for 3 h in the presence or absence of ACA (*n*=3). (**B**) Whole-cell patch clamp recording with 200 μM of cyclic ADP-ribose (cADPR) in pipette solution. At - 60 mV of holding voltage, development of inward current representing activation of TRPM2 by diffusion of cADPR to the cytosol was monitored in Mφs treated with carrier alone (BSA), PA alone or LPS alone for 4 h, or LP for a total of 4 h including LPS pretreatment for 3 h. Twenty μM of *N*[(*p*[amylcinnamoyl)anthranilic acid (ACA), a selective inhibitor of TRPM2, was applied to bath solution after confirming the steady-state activity inward current. (**C**) Amplitude of cADPR-induced inward current (b-c in B) (left group of the bar graph), and that of basal inward current inhibited by ACA (a-c in B) (right group of the bar graph) (*n*=19 for BSA; *n*=9 for PA; *n*=10 for LPS; *n*=9 for LP). (**D**) [Ca^2+^]_i_ in Mφs from *Trpm2*^+/+^ and *Trpm2*^-/-^ mice treated with LP for 1 h in the presence or absence of bafilomycin A1 emptying lysosomal Ca^2+^ reservoir after LPS pretreatment for 3 h, determined by ratiometric measurement after Fura-2 loading (*n*=10 for *Trpm2*^+/+^:BSA; *n*=27 for *Trpm2*^+/+^:LP; *n*=17 for *Trpm2*^+/+^:LP+BafA1; *n*=10 for *Trpm2*^-/-^:BSA; *n*=11 for *Trpm2*^-/-^:LP; *n*=9 for *Trpm2*^-/-^:LP+BafA1) (BafA1, bafilomycin A1). (**E**) Body weight of *Trpm2*^+/+^ and *Trpm2*^-/-^ mice on NCD (*n*=5) or HFD (*n*=8). (**F**) HOMA-IR index in *Trpm2*^+/+^ and *Trpm2*^-/-^ mice fed HFD for 8 weeks (*n*=5 for *Trpm2*^+/+^; *n*=8 for *Trpm2*^-/-^). Data shown as means ± SEM from more than 3 independent experiments. *p < 0.05, **p < 0.01 and ***p < 0.001 by one-way ANOVA with Tukey’s test (C, D, F) or two-way ANOVA with Tukey’s test (E).

**Figure 4−figure supplement 1.**
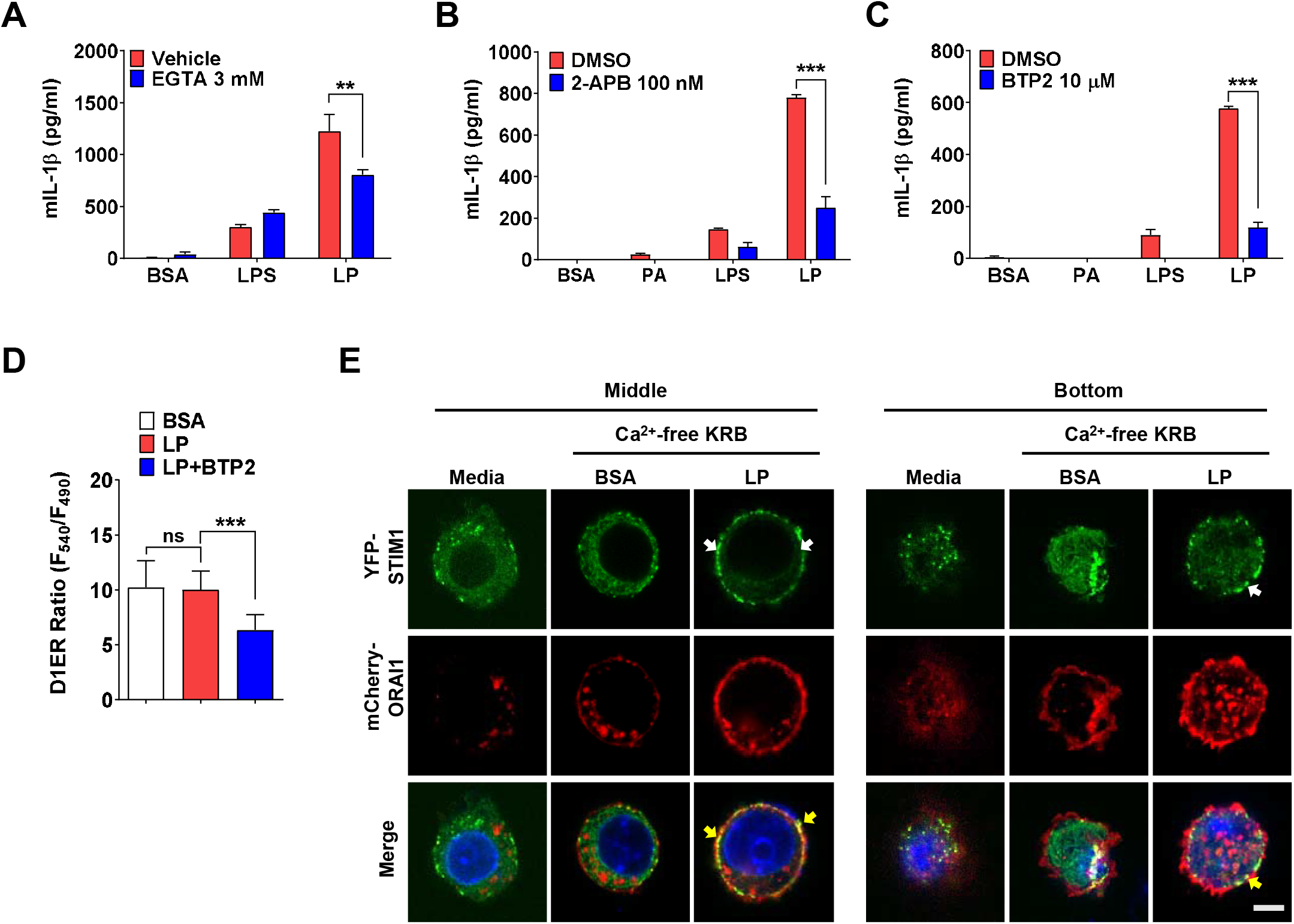
Store-operated Ca^2+^ entry (SOCE) in inflammasome by LP. (**A-C**) IL-1β ELISA of culture supernatant after treating Mφs with LPS alone or PA alone for 21 h, or with LP for a total of 21 h including LPS pretreatment for 3 h in the presence or absence of EGTA (A) (*n*=4), 2-APB (B) (*n*=4) or BTP2 (C) (*n*=3). (**D**) [Ca^2+^]_ER_ in *D1ER*-transfected BMDMs treated with LP for 1 h in the presence or absence of BTP2 without removal of extracellular Ca^2+^ after LPS pretreatment for 3 h (*n*=19 for BSA; *n*=16 for LP; *n*=24 for LP+BTP2). (**E**) STIM1 aggregation (white arrows) and colocalization with ORAI1 (yellow arrows) at the middle and bottom levels of BMDMs that were transfected with *YFP- STIM1* together with *mCherry-Orai1* and then treated with LP in Ca^2+^-free KRB buffer for 1 h after LPS pretreatment for 3 h, determined by confocal microscopy (scale bar, 5 mm). Data shown as means ± SEM from more than 3 independent experiments. **p < 0.01 and ***p < 0.001 by one-way ANOVA with Tukey’s test (D) or two-way ANOVA with Sidak test (A, B, C). Scale bar, 5 μm.

**Figure 5−figure supplement 1.**
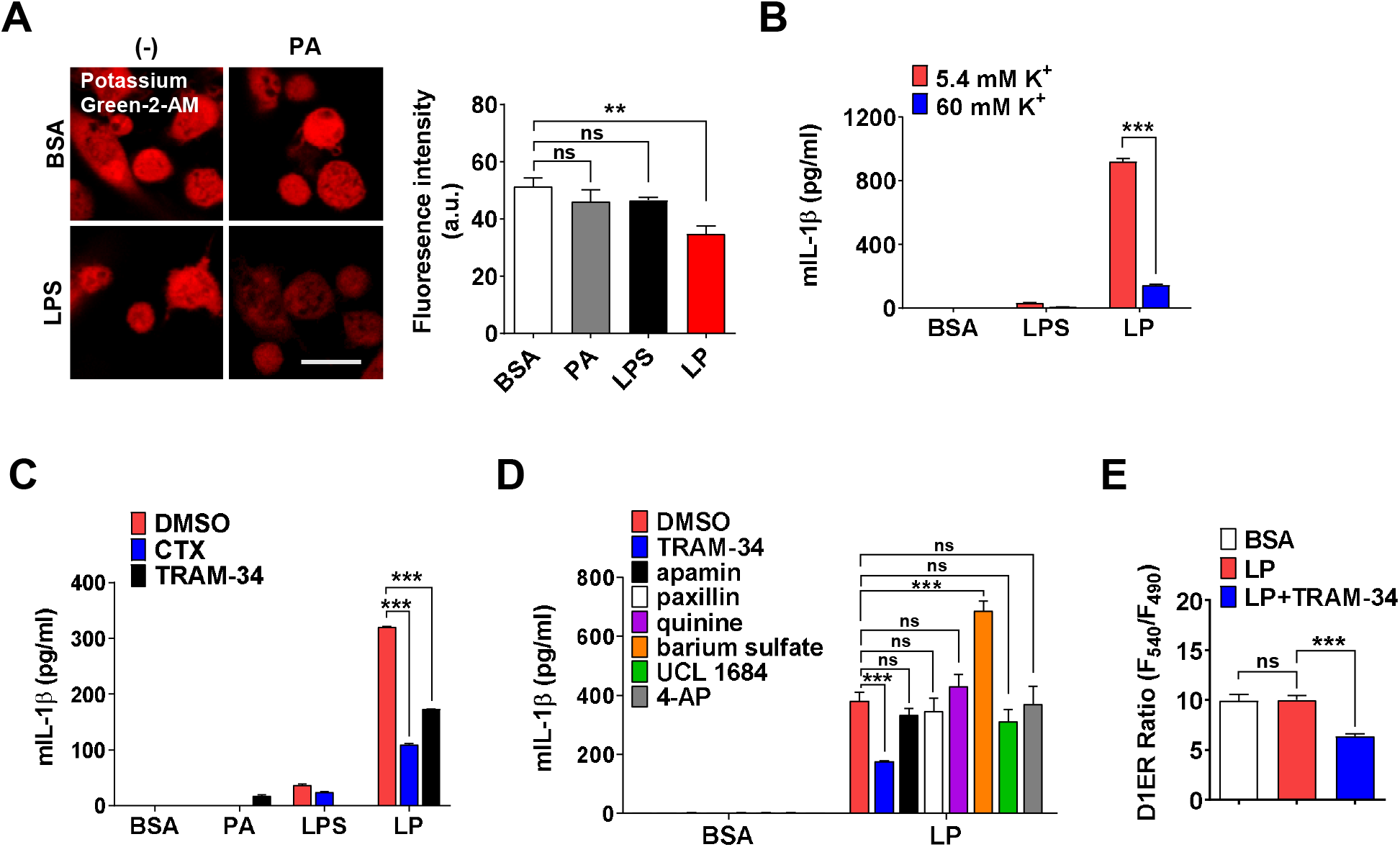
Effect of high extracellular K^+^ and inhibitors of K^+^ efflux channels on inflammasome. (**A**) [K^+^]_i_ in Mφs treated with LPS alone or PA alone for 21 h, or LP for a total of 21 h including LPS pretreatment for 3 h (right), determined by confocal microscopy after Potassium Green-2-AM loading. Representative Potassium Green-2 fluorescence images (left) (*n*=4 for BSA; *n*=2 for PA; *n*=4 for LPS; *n*=4 for LP). (**B**) IL-1β ELISA of culture supernatant after treating Mφs with LPS alone for 21 h or LP for a total of 21 h including LPS pretreatment for 3 h, at [K^+^]_e_ of 5.4 (K^+^ concentration in RPMI medium) or 60 mM (*n*=3 each). (**C, D**) IL-1β ELISA of culture supernatant after treating Mφs with LPS alone or PA alone for 21 h, or LP for a total of 21 h including LPS pretreatment for 3 h in the presence or absence of charybdotoxin (CTX) or TRAM-34 (C) (*n*=3), or several K^+^ efflux channel inhibitors (D) (*n*=3). (**E**) [Ca^2+^]_ER_ in *D1ER*-transfected BMDMs after LP treatment for 1 h in the presence or absence of TRAM-34 without extracellular Ca^2+^ removal after LPS pretreatment for 3 h (*n*=16). Data shown as means ± SEM from more than 3 independent experiments. **p < 0.01 and ***p < 0.001 by one-way ANOVA with Tukey’s test (A, E) or two-way ANOVA with Sidak test or with Tukey’s test (B, C, D). Scale bar, 20 μm.

**Figure 6−figure supplement 1.**
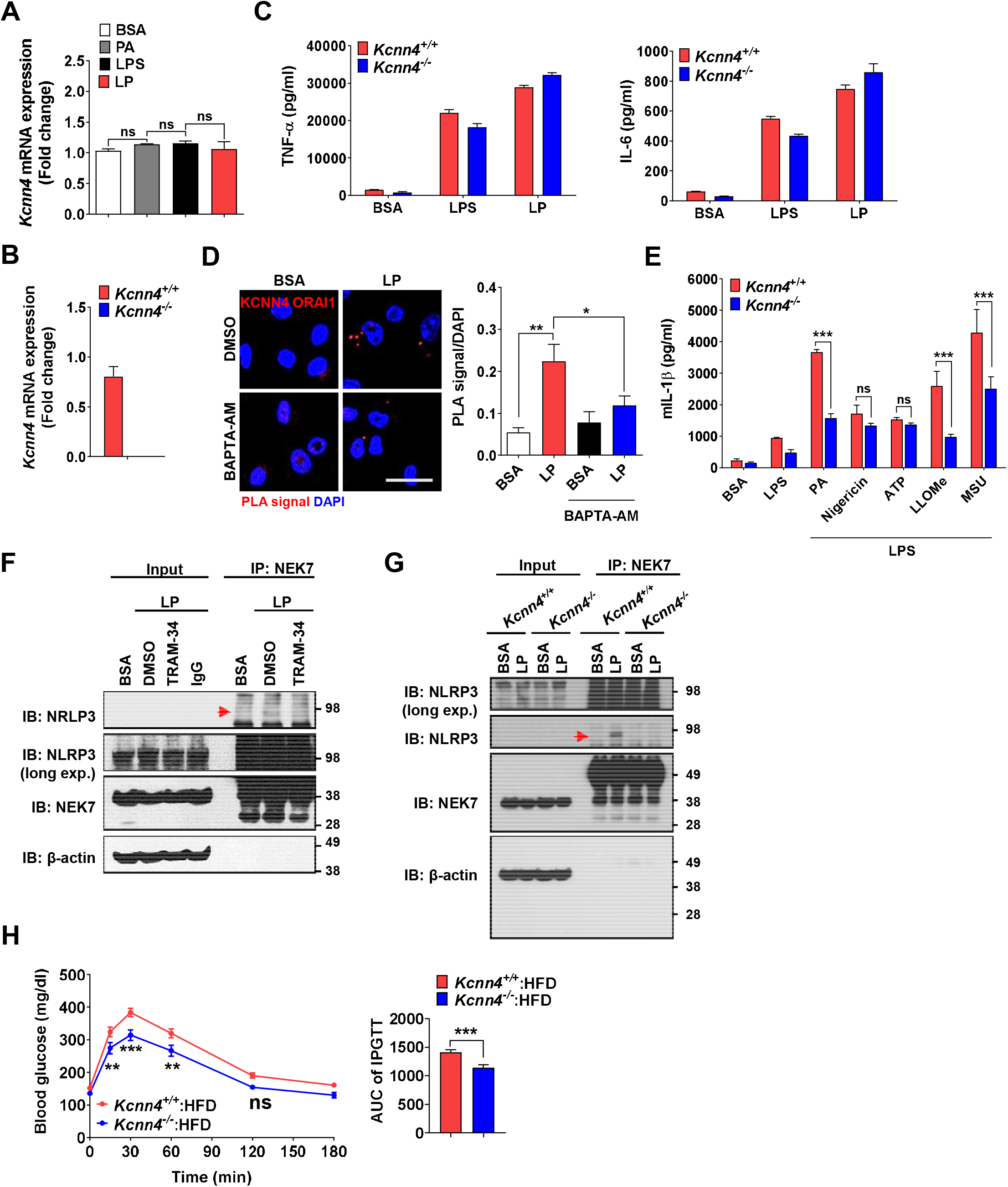
Effects of *Kcnn4* KO on metabolic profile and inflammasome. (**A**) Real-time RT-PCR of *Kcnn4* using mRNA from Mφs treated with PA alone or LPS alone for 21 h, or LP for a total of 21 h including LPS pretreatment for 3 h (*n*=3). (**B**) Real-time RT-PCR of *Kcnn4* using mRNA from Mφs of *Kcnn4*^+/+^ or *Kcnn4*^-/-^ mice (*n*=3). (**C**) TNF-α and IL-6 ELISA of culture supernatant after treating Mφs from *Kcnn4*^+/+^ or *Kcnn4*^-/-^ mice with LPS alone for 21 h or with LP for a total of 21 h including LPS pretreatment for 3 h (*n*=3). (**D**) PLA in Mφs treated with LP for a total of 21 h including LPS pretreatment for 3 h in the presence or absence of BAPTA-AM, using Abs specific for KCNN4 and ORAI1 (right). Representative fluorescence images (left) (scale bar, 20 μm) (*n*=10 for BSA; *n*=16 for LP; *n*=10 for BSA+BAPTA-AM; *n*=16 for LP+BAPTA-AM). (**E**) IL-1β ELISA of culture supernatant after treating Mφs from *Kcnn4*^+/+^ or *Kcnn4*^-/-^ mice with PA, nigericin, ATP, LLOMe and MSU for 18 h, 45 min, 1 h, 45 min and 3 h, respectively, after LPS pretreatment for 3 h (*n*=6). (**F**) Immunoprecipitation (IP) of Mφs treated with LP for a total of 21 h including LPS pretreatment for 3 h in the presence or absence of TRAM-34 using anti-NEK7 Ab and protein G beads. After bead heating in a sample buffer, collected supernatant was subjected to immunoblotting (IB) using indicated Abs. (red arrow, band of the correct size) (**G**) Immunoprecipitation of Mφs from *Kcnn4*^+/+^ or *Kcnn4*^-/-^ mice treated with LP for a total of 21 h including LPS pretreatment for 3 h using anti-NEK7 Ab and protein G beads. After bead heating in a sample buffer, collected supernatant was subjected to IB using indicated Abs. (red arrow, band of the correct size) (**H**) Intraperitoneal glucose tolerance test (IPGTT) in *Kcnn4*^+/+^ and *Kcnn4*^-/-^ mice fed HFD for 8 weeks (left). AUC (right) (*n*=9). Data shown as means ± SEM from more than 3 independent experiments. *p < 0.05, **p < 0.01 and ***p < 0.001 by one-way ANOVA with Tukey’s test (A, D), two-tailed Student’s *t*-test (B, H), or two-way ANOVA with Sidak test (C, E).

**Figure 7−figure supplements 1.**
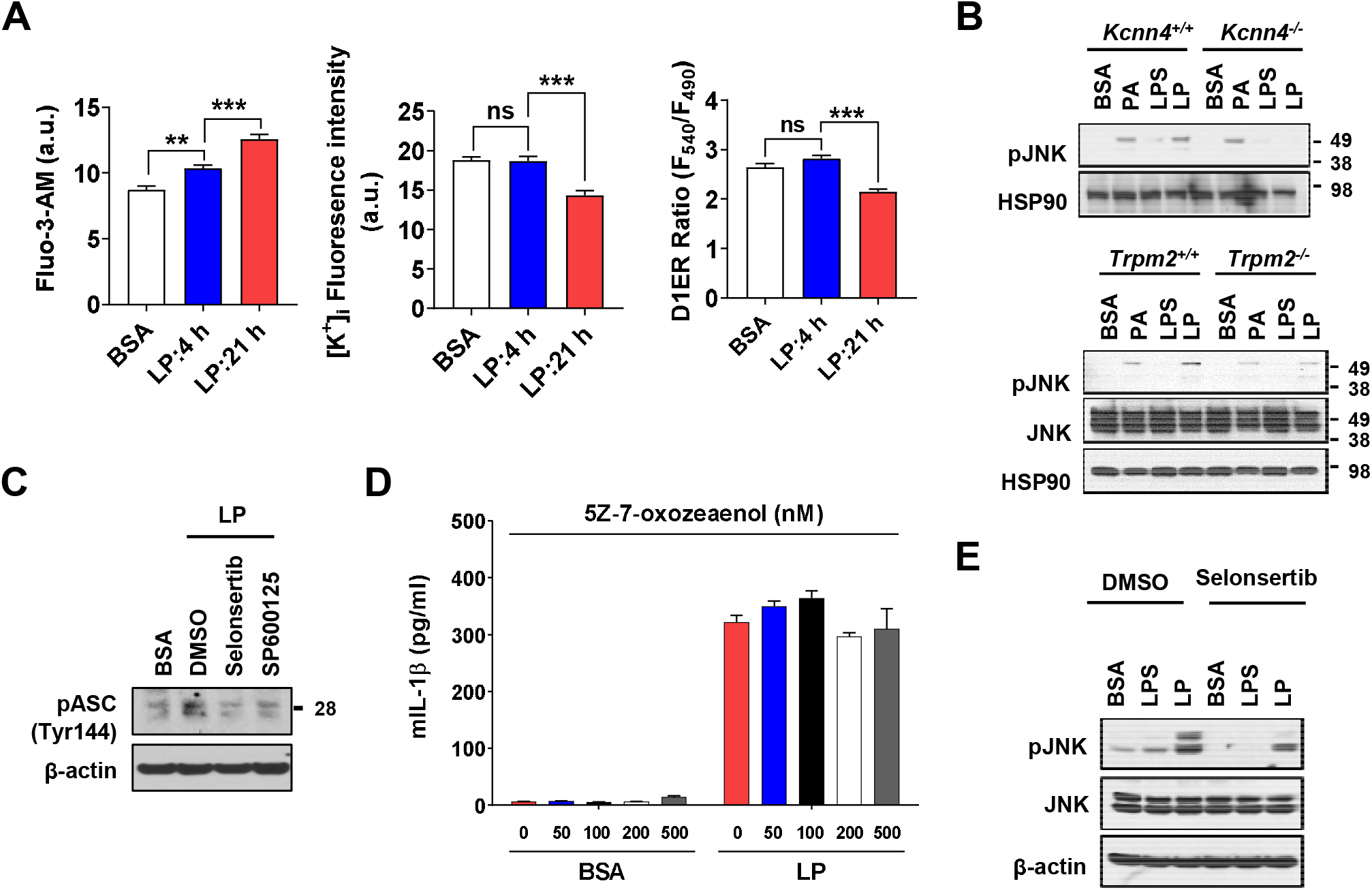
Effect of *Kcnn4* KO and inhibitor of TAK1 or ASK1 on activation of inflammasome and JNK. (**A**) [Ca^2+^]_i_ (left), [K^+^]_i_ (middle) and [Ca^2+^]_ER_ (right) determined by Flou-3-AM staining, Potassium Green-2-AM staining and *D1ER* transfection, respectively, after treatment with LP for a total of 4 or 21 h including LPS pretreatment for 3 h (*n*=10 for Fluo-3-AM; *n*=10 for Potassium Green-2-AM; *n*=50 for *D1ER*). (**B**) IB of lysate of Mφs from *Kcnn4*^+/+^ and *Kcnn4*^-/-^ mice (left) or *Trpm2*^+/+^ and *Trpm2*^-/-^ mice (right) treated with PA alone for 21 h, LPS alone for 21 h or LP for a total of 21 h including LPS pretreatment for 3 h, using indicated Abs. (**C**) IB of lysate of BMDMs treated with LP for a total of 21 h including LPS pretreatment for 3 h in the presence or absence of ASK1 (selonsertib) or JNK inhibitor (SP600125) using indicated Ab. (**D**) IL-1β ELISA of culture supernatant after treating BMDMs with LP for a total of 21 h including LPS pretreatment for 3 h in the presence or absence of 5Z-7-oxozeaenol (*n*=3). (**E**) IB of lysate of BMDMs treated with LPS alone for 21 h or with LP for a total of 21 h including LPS pretreatment for 3 h in the presence or absence of selonsertib using indicated Abs. Data shown as means ± SEM from more than 3 independent experiments*p < 0.05, **p < 0.01 and ***p < 0.001 by one-way ANOVA with Tukey’s test (A), or two-way ANOVA with Tukey’s test (D).

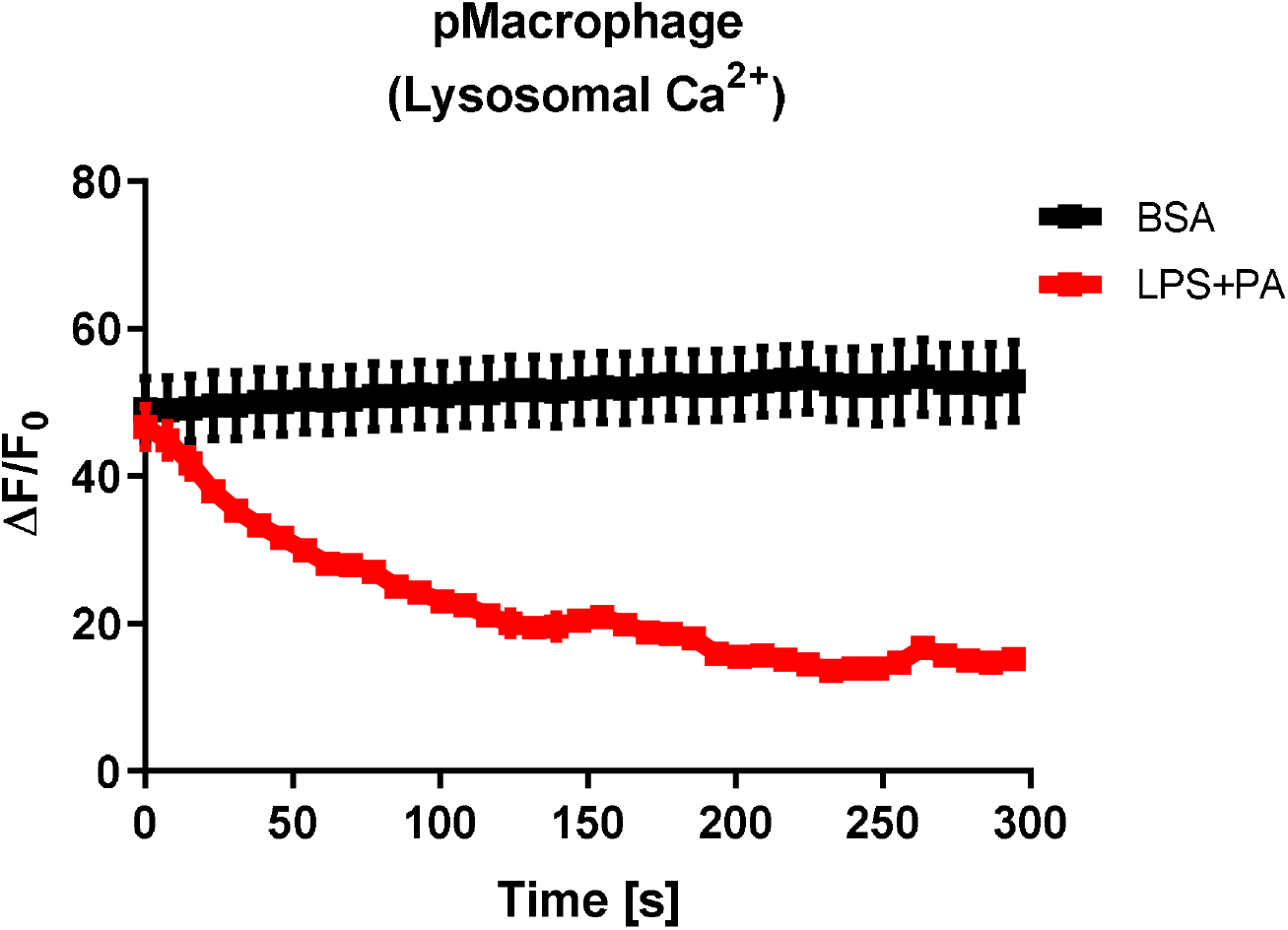

## References

Agarwal JJ, Zhu Y, Zhang Q-Y, Mongin AA, Hough LB 2014. TRAM-34, a putatively selective blocker of intermediate-conductance, calcium-activated potassium channels, inhibits cytochrome P450 activity. PLOS ONE 8, e63028.

Akaike N, Harata N 1994. Nystatin perforated patch recording and its applications to analyses of intracellular mechanisms. Japanese Journal of Physiology 44, 433–473.

An D, Hao F, Hu C, Kong W, Xu X, Cui M-Z 2018. JNK1 Mediates Lipopolysaccharide-Induced CD14 and SR-AI Expression and Macrophage Foam Cell Formation. Frontiers in Physiology 8, 1705.

Bachar E, Ariav Y, Ketzinel-Gilad M, Cerasi E, Kaiser N, Leibowitz G 2009. Glucose amplifies fatty acid-induced endoplasmic reticulum stress in pancreatic beta-cells via activation of mTORC1. PLOS ONE 4, e4954.

Bock KW 2020. Aryl hydrocarbon receptor (AHR) functions: Balancing opposing processes including inflammatory reactions. Biochemical Pharmacology 178, 114093.

Bogeski I, Al-Ansary D, Qu B, Niemeyer BA, Hoth M, Peinelt C 2010. Pharmacology of ORAI channels as a tool to understand their physiological functions. Expert Review of Clinical Pharmacology 3, 291–303.

Campden RI, Zhang Y 2019. The role of lysosomal cysteine cathepsins in NLRP3 inflammasome activation Archives of Biochemistry and Biophysics 670, 32–42.

Chen M, Fei Y, Chen T-Z, Li Y-G, Chen P-S 2021. The regulation of the small-conductance calcium- activated potassium current and the mechanisms of sex dimorphism in J wave syndrome. Pflugers Archive 473, 492–506.

Chen R, Chung S-H 2013. Molecular dynamics simulations of scorpion toxin recognition by the Ca(2+)-activated potassium channel KCa3.1. Biophysical Journal 105, 1829–1837.

Derler I, Jardin I, Romanin C 2016. Molecular mechanisms of STIM/Orai communication. American Journal of Physiology 310, C643–662.

Di A, Xiong S, Ye Z, Malireddi RKS, Kometani S, Zhong M, Mittal M, Hong Z, Kanneganti TV, Rehman J, Malik AB 2017. The TWIK2 potassium influx channel in macrophages mediates NLRP3 inflammasome-induced inflammation. Immunity 49, 56–65.

Drinkall S, Lawrence CB, Ossola B, Russell S, Bender C, Brice NB, Dawson LA, Harte M, Brough D 2022. The two pore potassium channel THIK-1 regulates NLRP3 inflammasome activation. Glia 70, 1301–1316.

Duffy SM, Ashmole I, Smallwood DT, Leyland ML, Bradding P 2015. Orai/CRACM1 and KCa3.1 ion channels interact in the human lung mast cell plasma membrane. Cell Communcation and Signaling 13, 32.

Ferreira R, Schlichter LC 2013. Selective activation of KCa3.1 and CRAC channels by P2Y2 receptors promotes Ca(2+) signaling, store refilling and migration of rat microglial cells PLOS ONE 8, e62345.

Gao Y-d, Hanley PJ, Rinné S, Zuzarte M, Daut J 2010. Calcium-activated K(+) channel (K(Ca)3.1) activity during Ca(2+) store depletion and store-operated Ca(2+) entry in human macrophages. Cell Calcium 48, 19–27.

Garrity AG, Wang W, COllier CMD, Levery SA, Gao Q, Xu H 2016. The endoplasmic reticulum, not the pH gradient, drives calcium refilling of lysosomes. eLife 5, e15887.

González C, Baez-Nieto D, Valencia I, Oyarzún I, Rojas P, Naranjo D, Latorre R 2012. K(+) channels: function-structural overview. Comprehensive Physiology 2, 2087–2149.

Groß C, Mishra R, Schneider KS, Médard G, Wettmarshausen J, Dittlein DC, Shi H, Gorka O, Koenig PA, Fromm S, et al. 2016. K+ efflux-independent NLRP3inflammasome activation by small molecules targeting mitochondria. Immunity 45, 761–773.

Grynkiewicz G, Poenie M, Tsien RY 1985. A new generation of Ca2+ indicators with greatly improved fluorescence properties. Journal of Biological Chemistry 260, 3440–3450.

Guéguinou M, Chantôme A, Fromont G, Bougnoux P, Vandier C, Potier-Cartereau M 2014. KCa and Ca2+channels: The complex though. Biochimica et Biophysica Acta 1843, 2322–2333.

Hara H, Tsuchiya K, Kawamura I, Fang R, Hernandez-Cuellar E, Shen Y, Mizuguchi J, Schweighoffer E, Tybulewicz V, Mitsuyama M 2013. Phosphorylation of the adaptor ASC acts as a molecular switch that controls the formation of speck-like aggregates and inflammasome activity. Nature Immunology 14, 1247–1255.

He Y, Zeng MY, Yang D, Motro B, Núñez G 2016. NEK7 is an essential mediator of NLRP3 activation downstream of potassium efflux. Nature 530, 354–357.

Hofer AM, Fasolato C, Pozzan T 1998. Capacitative Ca2+ entry is closely linked to the filling state of internal Ca2+ stores: a study using simultaneous measurements of ICRAC and Intraluminal [Ca2+]. Journal of Cell Biology 130, 325–334.

Hornung V, Bauernfeind F, Halle A, Samstad EO, H. K, Rock KL, Fitzgerald KA, Latz E 2008. Silica crystals and aluminum salts activate the NALP3 inflammasome through phagosomal destabilization. Nature Immunology 9, 847–856.

Immanuel CN, Teng B, Dong B, Gordon EM, Kennedy JA, Luellen C, Schwingshackl A, Cormier SA, Fitzpatrick EA, Waters CM 2019. Apoptosis Signal-Regulating kinase-1 Promotes Inflammasome Priming in Macrophages. American Journal of Physiology 316, L418–427.

Katsnelson MA, Rucker LG, Russo HM, Dubyak GR 2015. K+ efflux agonists induce NLRP3 inflammasome activation independently of Ca2+ signaling. Journal of Immunology 194, 3937–3952.

Kinnear NP, Boittin F-X, Thomas JM, Galione A, Evan AM 2004. Lysosome-sarcoplasmic reticulum junctions. A trigger zone for calcium signaling by nicotinic acid adenine dinucleotide phosphate and endothelin-1. Journal of Biological Chemistry 279, 54319–54326.

Korge P, Honda HM, Weiss JN 2003. Effects of fatty acids in isolated mitochondria: implications for ischemic injury and cardioprotection. American Journal of Physiology 285, H295–H269.

Kraft R, Grimm C, Frenzel H, Harteneck C 2006. Inhibition of TRPM2 cation channels by N-(p-amylcinnamoyl)anthranilic acid. British Journal of Pharmacology 148, 264–273.

Lange I, Yamamoto S, Partida-Sanchez S, Mori Y, Fleig A, Penner R 2009. TRPM2 functions as a lysosomal Ca2+-release channel in beta cells. Science Signaling 2.

Lee GS, Subramanian N, Kim A, Aksentijevich I, Goldbach-Mansky R, Sacks DB, Germain RN, Kastner DL, Chae JJ 2012. The calcium-sensing receptor regulates the NLRP3 inflammasome through Ca2+ and cAMP. Nature 492, 123–127.

Lee H-Y, Kim J, Quan Y, Lee J-C, Kim M-S, Km S, Bae J-W, Hur KY, Lee MS 2016. Autophagy deficiency in myeloid cells increases susceptibility to obesity-induced diabetes and experimental colitis. Autophagy 12, 1390–1403.

Li Y, Chen M, Xu Y, Yu X, Xiong T, Du M, Sun J, Liu L, Tang Y, Yao P 2016. Iron-mediated lysosomal membrane permeabilization in ethanol-induced hepatic oxidative damage and apoptosis: protective effects of quercetin. Oxidative Medicine and Cellular Longevity 2016, 4147610.

Madry C, Kyrargyri V, Arancibia-Cárcamo IL, Renaud Jolivet R, Kohsaka S, Bryan RM, Attwell D 2018. Microglial ramification, surveillance, and interleukin-1β release are regulated by the two-pore domain K + channel THIK-1. Neuron 97, 299–312.

Maggi CA, Santicioli P, Geppetti P, Parlani M, Astolfi M, Del Bianco E, Patacchini R, Giuliani S, Meli A 1989. The effect of calcium free medium and nifedipine on the release of substance P-like immunoreactivity and contractions induced by capsaicin in the isolated guinea-pig and rat bladder. General Pharmacology 20, 445–456.

Misawa T, Takahama M, Kozaki T, Lee H, Zou J, Saitoh T, Akira S 2013. Microtubule-driven spatial arrangement of mitochondria promotes activation of the NLRP3 inflammasome. Nature Immunology 14, 454–460.

Muñoz-Planillo R, Kuffa P, Martínez-Colón G, Smith BL, Rajendiran TM, Núñez G 2013. K+ efflux is the common trigger of NLRP3 inflammasome activation by bacterial toxins and particulate matter. Immunity 38, 1142–1153.

Nakamura S, Takamura T, Matsuzawa-Nagata N, Takayama H, Misu H, Noda H, Nabemoto S, Kurita S, Ota T, Ando H, et al. 2009. Palmitate induces insulin resistance in H4IIEC3 hepatocytes through reactive oxygen species produced by mitochondria. Journal of Biological Chemistry 284, 14809–14818.

Nam T-S, Park D-R, Rah S-Y, Woo T-G, Chung H-T, Brenner C, Kim U-H 2020. Interleukin-8 drives CD38 to form NAADP from NADP + and NAAD in the endolysosomes to mobilize Ca 2+ and effect cell migration. FASEB Journal 34, 12565–12576.

Okada M, Matsuzawa A, Yoshimura A, Ichijo H 2014. The lysosome rupture-activated TAK1-JNK pathway regulates NLRP3 inflammasome activation. Journal of Biological Chemistry 289, 32926–32936.

Park K, Lim H, Kim J, Hwang H, Lee YS, Bae SH, Kim H, Kim H, Kang S-W, Kim JW, Lee M-S 2022. Essential role of lysosomal Ca2+-mediated TFEB activation in mitophagy and functional adaptation of pancreatic ΣΙ-cells to metabolic stress. Nature Communications 13, 1300.

Park KS, Poburko D, Wollheim CB, Demaurex N 2009. Amiloride derivatives induce apoptosis by depleting ER Ca(2+) stores in vascular endothelial cells. British Journal of Pharmacology 156, 1296–1304.

Penny CJ, Kilpatrick BS, Han JM, Sneyd J, Patel S 2014. A computational model of lysosome-ER Ca2+ microdomains. Journal of Cell Science 127, 2934–2943.

Place DE, Sami P, Karki R, Briard B, Vogel P, Kanneganti TD 2018. ASK family kinases are required for optimal NLRP3 inflammasome priming. American Journal of Pathology 188, 1021–1030.

Raffaello A, Mammucari C, Gherardi G, Rizzuto R 2016. Calcium at the center of cell signaling: interplay between endoplasmic reticulum, mitochondria, and lysosomes. Trends in Biochemical Sciences 41, 1035–1049.

Sanurjo CIL, Tovey SC, Taylor CW 2014. Rapid recycling of Ca2+ between IP3-sensitive stores and lysosomes. PLOS ONE 9, e111275.

Schoenmakers TJ, Visser GJ, Flik G, Theuvenet AP 1992. CHELATOR: an improved method for computing metal ion concentrations in physiological solutions. Biotechniques 12, 870–874.

Schroeder ME, Russo S, Costa C, Hori J, Tiscornia I, Bollati-Fogolín M, Zamboni DS, Ferreira G, Cairoli E, Hill M 2017. Pro-inflammatory Ca++-activated K+ channels are inhibited by hydroxychloroquine. Scientific Reports 7, 1892.

Sharif H, Wang L, Wang WL, Magupalli VG, Andreeva L, Qiao Q, Hauenstein AV, Wu Z, Núñez G, Mao Y, Wu H 2019. Structural mechanism for NEK7-licensed activation of NLRP3 inflammasome. Nature 570, 338–343.

Shen D, Wang X, Li X, Zhang X, Yao Z, Dibble S, Dong XP, Yu T, Lieberman AP, Showalter HD, Xu H 2012. Lipid storage disorders block lysosomal trafficking by inhibiting a TRP channel and lysosomal calcium release. Nature Communications 3, 731.

Strøbæk D, Jørgensen TD, Christophersen P, Ahring PK, Olesen S-P 2000. Pharmacological characterization of small-conductance Ca2+-activated K+ channels stably expressed in HEK 293 cells. British Journal of Pharmacology 129, 991–999.

Sumoza-Toledo A, Penner R 2010. TRPM2: a multifunctional ion channel for calcium signalling. Journal of Physiology 589.7, 1515–1525.

Suzuki J, Kanemaru K, Ishii K, Ohkura M, Lino M 2014. Imaging intraorganellar Ca2+ at subcellular resolution using CEPIA. Nature Communications 5, 4153.

Tahtinen S, Tong A-J, Himmels P, Oh J, Paler-Martinez A, Kim L, Wichner S, Oei Y, McCarron MJ, Freund EC, et al. 2022. IL-1 and IL-1ra are key regulators of the inflammatory response to RNA vaccines. Nature Immunology 23, 532–542.

Tardiolo G, Bramanti P, Mazzon E 2018. Overview on the effects of N-acetylcysteine in neurodegenerative diseases. Molecules 23, 3305.

Tseng HSL, Vong CT, Kwan YW, Lee SM-Y, Hoi MPM 2017. TRPM2 regulates TXNIP-mediated NLRP3 inflammasome activation via interaction with p47 phox under high glucose in human monocytic cells. Scientific Reports 6, 35016.

Uchida K, Tominaga M 2011. TRPM2 modulates insulin secretion in pancreatic b-cells’. Islets 3, 209–211.

Vaca L 2010. SOCIC: the store-operated calcium influx complex. Cell Calcium 47, 199–209.

Wang L, Negro R, Wu H 2020. TRPM2, linking oxidative stress and Ca2+ permeation to NLRP3 inflammasome activation. Current Opinion in Immunology 62, 131–135.

Weber K, Schilling JD 2014. Lysosomes integrate metabolic-inflammatory cross-talk in primary macrophage inflammasome activation. Journal of Biological Chemistry 289, 9158–9171.

Weisberg SP, McCann D, Desai M, Rosenbaum M, Leibel RL, Ferrante AWJ 2003. Obesity is associated with macrophage accumulation in adipose tissue. Journal of Clinical Investigation 112, 1797–1808.

Wen H, Gris D, Lei Y, Jha S, Zhang L, Huang MT, Brickey WJ, Ting JP 2011. Fatty acid-induced NLRP3-ASC inflammasome activation interferes with insulin signaling. Nature Immunology 12, 408–415.

Wulff H, Miller MJ, Hänsel W, Grissmer S, Cahalan MD, Chandy KG 2000. Design of a potent and selective inhibitor of the intermediate-conductance Ca2+-activated K+ channel, IKCa1: A potential immunosuppressant. Proceedings of the National Academy of Science USA 97, 8156–8160.

Xian H, Watari K, Sanchez-Lopez E, Offenberger J, Onyuru J, Sampath H, Ying W, Hoffman HM, Shadel GS, Karin M 2022. Oxidized DNA fragments exit mitochondria via mPTP- and VDAC- dependent channels to activate NLRP3 inflammasome and interferon signaling. Immunity in press.

Xu S, Nam SM, Kim JH, Das R, Choi SK, Nguyen TT, Quan X, Choi SJ, Chung CH, Lee EY, et al. 2015. Palmitate induces ER calcium depletion and apoptosis in mouse podocytes subsequent to mitochondrial oxidative stress. Cell Death and Disease 6, e1976.

Xu Z, Chen Z-M, Wu X, Zhang L, Cao Y, Zhou P 2020. Distinct molecular mechanisms underlying potassium efflux for NLRP3 inflammasome activation. Frontiers in Immunology 11, 609441.

Yang D, He Y, Muñoz-Planillo R, Liu Q, Núñez G 2015. Caspase-11 Requires the Pannexin-1 Channel and the Purinergic P2X7 Pore to Mediate Pyroptosis and Endotoxic Shock. Immunity 43, 923–932.

Yang J, Zhao Z, Gu M, Feng X, Xu H 2019. Release and uptake mechanisms of vesicular Ca2+ store. Protein & Cell 10, 8–19.

Yaron JR, Gangaraju S, Rao MY, Kong X, Zhang L, Su F, Tian Y, Glenn HL, Meldrum DR 2015. K(+) regulates Ca(2+) to drive inflammasome signaling: dynamic visualization of ion flux in live cells. Cell Death and Disease 6, e1964.

Zhang X, Cheng X, Yu L, Yang. J,., Calvo R, Patnaik S, Hu X, Gao Q, Yang M, Lawas M, et al. 2016. MCOLN1 is a ROS sensor in lysosomes that regulates autophagy. Nature Communications 7, 12109.

Zhang Y, Yang Y, Yu H, Li M, Hang L, Xu X 2020. Apigenin protects mouse retina against oxidative damage by regulating the Nrf2 pathway and autophagy Oxidative Medicine and Cellular Longevity 2020, 9420704.

Zhang Z, Zhang W, Jung DY, Ko HJ, Lee Y, Friedline RH, Lee E, Jun J, Ma Z, Kim F, et al. 2012. TRPM2 Ca2+ channel regulates energy balance and glucose metabolism. American Journal of Physiology 302, E807–816.

Zhong Z, Zhai Y, Liang S, Mori Y, Han R, Sutterwala FS, Qiao L 2013. TRPM2 links oxidative stress to NLRP3 inflammasome activation. Nature Communications 4, 1611.

Zhou R, Yazdi AS, Menu P, Tshopp J 2011. A role for mitochondria in NLRP3 inflammasome activation. Nature 469, 221–226.

Zitt C, Strauss B, Schwarz EC, Spaeth N, Rast G, Hatzelmann A, Hoth M 2004. Potent inhibition of Ca2+ release-activated Ca2+ channels and T-lymphocyte activation by the pyrazole derivative BTP2. Journal of Biological Chemistry 279, 12427–12437.

